# Functional proximity mapping of RNA binding proteins uncovers a mitochondrial mRNA anchor that promotes stress recovery

**DOI:** 10.1101/2020.11.17.387209

**Authors:** Wei Qin, Samuel A Myers, Dominique K. Carey, Steven A Carr, Alice Y Ting

**Affiliations:** Departments of Biology, Genetics, and Chemistry, Stanford University, Stanford, United States; The Broad Institute of MIT and Harvard, Cambridge, Massachusetts, United States

## Abstract

Proximity labeling (PL) with genetically-targeted promiscuous enzymes has emerged as a powerful tool for unbiased proteome discovery. By combining the spatiotemporal specificity of PL with methods for functional protein enrichment, it should be possible to map specific protein subclasses within distinct compartments of living cells. Here we demonstrate this capability for RNA binding proteins (RBPs), by combining peroxidase-based PL with organic-aqueous phase separation of crosslinked protein-RNA complexes (“APEX-PS”). We validated APEX-PS by mapping nuclear RBPs, then applied it to uncover the RBPomes of two unpurifiable subcompartments - the nucleolus and the outer mitochondrial membrane (OMM). At the OMM, we discovered the RBP SYNJ2BP, which retains specific nuclear-encoded mitochondrial mRNAs during translation stress, to promote their local translation and import of protein products into the mitochondrion during stress recovery. APEX-PS is a versatile tool for compartment-specific RBP discovery and expands the scope of PL to *functional* protein mapping.

**Graphic Abstract:** 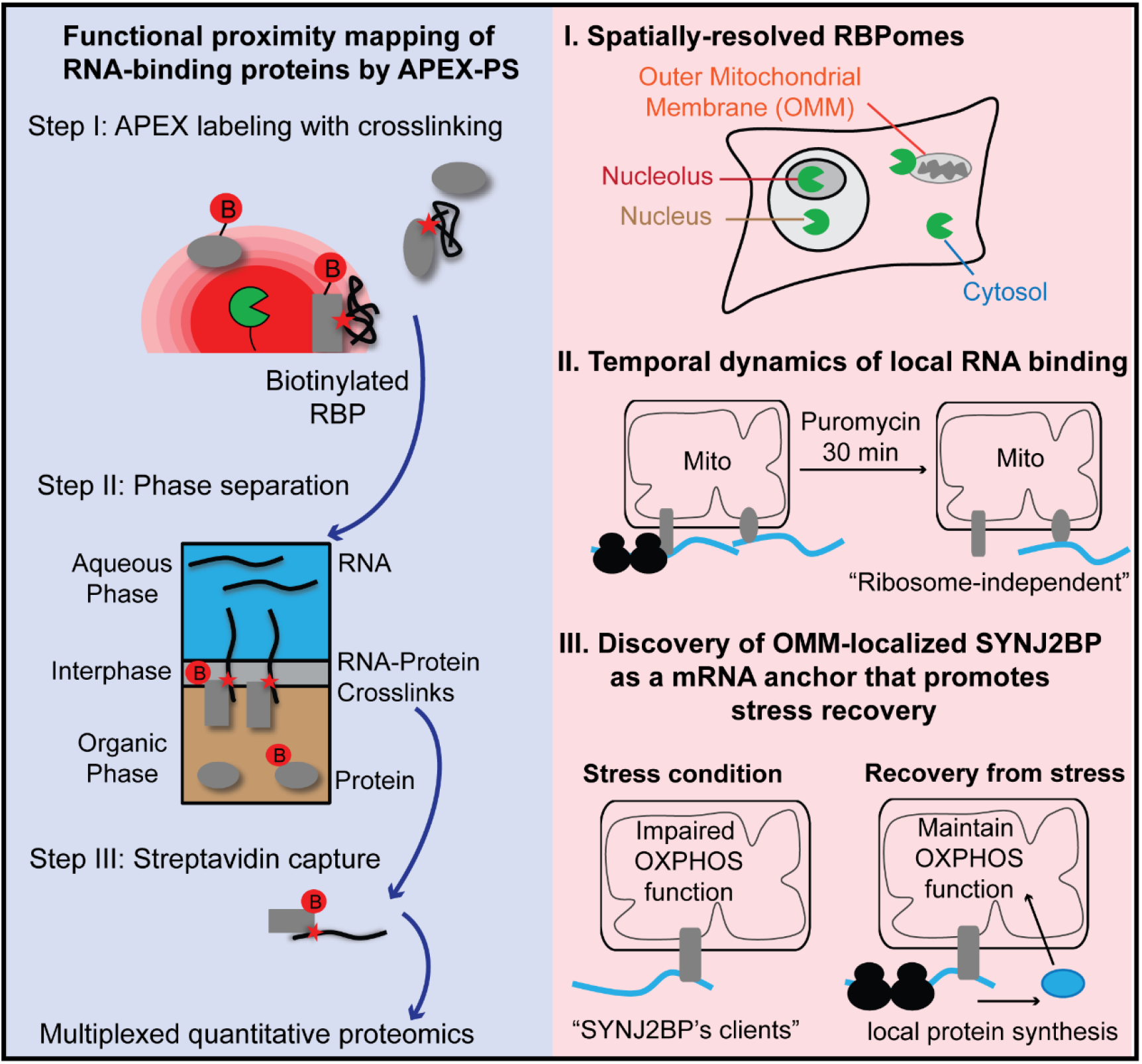

## Introduction

RNA-protein interactions are pervasive in both transient and stable macromolecular complexes underlying transcription, translation and stress response ^1, 2^. Consequently, RNA binding proteins (RBPs) constitute an important and large (estimated ∼10%) functional subclass of the human proteome ^3-5^. While methods for the global discovery of RBPs, such as oligo(dT) bead capture ^6-9^, metabolic RNA labeling ^10, 11^, and organic-aqueous phase separation ^12-14^, are highly valuable, a richer understanding of RBP function could be achieved with spatially- and temporally-resolved approaches that report not only the proteins that have RNA binding function but where they are located within the cell and under what conditions. For instance, RBPs in the nucleolus are candidates for ribosome biogenesis and assembly, while RBPs at the outer mitochondrial membrane (OMM) are candidates for mRNA recruitment for local translation of mitochondrial proteins. RBPs in mitochondrial RNA granules may participate in gene expression, which are dynamically regulated by mitochondrial fission and fusion^15^. A spatiotemporally-resolved approach for RBP discovery could yield valuable insights into organelle- or region-specific questions concerning the dynamics of RNA-protein interactions.

To develop such a method, we sought to combine the spatial proteomics method known as “proximity labeling” (PL) with a method for functional RBP enrichment. In PL, an engineered promiscuous labeling enzyme such as APEX^16^, TurboID^17^, or BioID^18^ is targeted via genetic fusion to a specific subcellular compartment or macromolecular complex. Addition of a small molecule substrate to live cells results in catalytic generation of a reactive biotin species (such as biotin-phenoxyl radical or biotin-AMP) from the active site of the promiscuous enzyme, which diffuses outward to covalently tag proximal endogenous proteins. PL methods have been used to map organelle proteomes (mitochondrion, ER, lipid droplets, stress granules)^19^, dynamic interactomes^20, 21^, and in vivo secretomes^22, 23^. Recently, PL has been extended to spatial mapping of transcriptomes in living cells ^24-26^.

While PL is a powerful technology for *unbiased* discovery of proteins or RNA in specific subcellular locales, in many cases one wishes to focus a search on a specific protein or RNA functional subclass. PL has not previously been used in the manner of activity-based protein profiling^27, 28^, for instance, to enrich specific enzyme families.

If one could combine the spatiotemporal specificity and live cell compatibility of PL with targeted approaches that enrich proteins or RNA based on functional activity (“functional proximity labeling”), it would enable the simultaneous assignment of localization, timing, and function to specific protein populations.

Here, we develop such methodology and apply it to study RBPs in relation to mitochondria and stress. RBPs play an important role in cellular responses to environmental stresses, including heat stress, oxidative pressure and nutrient deprivation. Under stress conditions, protein translation is severely impaired and translationally-stalled mRNAs rapidly bind to cytosolic RBPs, driving the formation of stress granules to promote cell survival and recovery^29^. As cells recover from stress, disassembly of stress granules proceeds in a stepwise manner, and specific subsets of RNAs are expected to be primed for re-entry into translation^30^. This current understanding of the stress response raises the question of how cells handle essential transcripts which may need to re-enter translation immediately upon stress recovery. In particular, specific mitochondrial transcripts may need to be immediately available for translation in order to enable energy production and restoration of cellular homeostasis. Are there specific mechanisms to protect essential mitochondrial transcripts such that they do not become trapped in stress granules, and are instead immediately available for translation upon stress recovery?

## Results

### Design and validation of APEX-PS for functional proximity labeling of subcellular RBPs

To develop an approach for tandem PL and functional RBP enrichment, we first selected the fastest PL enzyme available - APEX2 - an engineered peroxidase that catalyzes the one-electron oxidation of biotin-phenol conjugates^31^. The radical product of this oxidation forms covalent adducts with electron rich protein sidechains such as tyrosine and has a half-life of <1 millisecond, resulting in labeling radii <10 nanometers. Labeling is initiated by the addition of biotin-phenol and H_2_O_2_ to live cells and typically performed within 20 seconds to 1 minute.

To adapt APEX-based PL for spatiotemporally-resolved enrichment of RBPs specifically, we envisioned following the 1-minute live cell APEX biotinylation with RNA-protein crosslinking using formaldehyde (FA) or UV (**Fig. 1a**), then enriching crosslinked RNA-protein complexes via organic-aqueous phase separation (PS). Previous studies have shown that while proteins partition to the organic layer and RNAs partition to the aqueous layer, crosslinked RNA-protein complexes partition to the interphase (**Fig. 1b**) and can be enriched from this layer. The OOPS and XRNAX methods^12, 13^ used this approach to identify RBPs on a large-scale, albeit with loss of spatial information. Following phase separation (PS) to enrich RBPs, we perform streptavidin-based enrichment of APEX-biotinylated proteins, as in our standard PL workflow. The intersection of APEX-based PL with PS (“APEX-PS”) should enable the compartment-specific identification of RBPs with minutes-long temporal resolution.

**Fig. 1.**
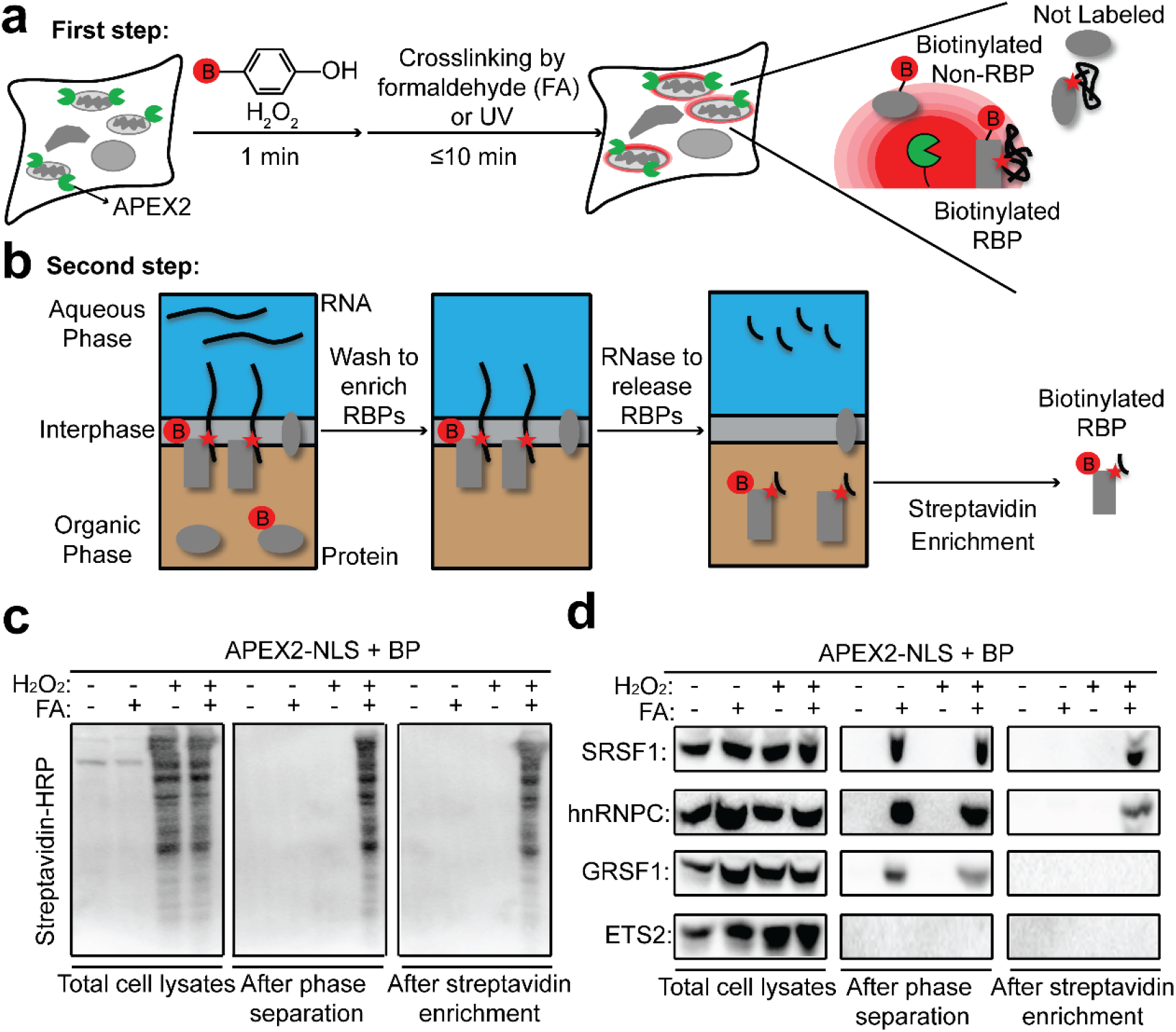
Development and validation of APEX-PS. **a**, First step: 1-minute APEX-catalyzed biotinylation in live cells followed by RNA-protein crosslinking. Red B, biotin. RBP, RNA binding protein. **b**, Second step: enrichment of biotinylated RBPs by phase separation and streptavidin bead-based purification. Crosslinked protein-RNA complexes localize to the interphase, while free proteins and RNA localize to the organic phase and aqueous phase, respectively. After washing, RNA-crosslinked proteins are released by RNase treatment and subjected to streptavidin bead-based enrichment. **c**, Streptavidin blotting reveals enrichment of nuclear RBPs by APEX-PS-NLS. Biotinylation and formaldehyde crosslinking was performed in HEK293T cells expressing nuclear-localized APEX (APEX-NLS). Streptavidin blotting was performed on total cell lysate (left), samples after phase separation (middle), and samples after both phase separation and streptavidin enrichment (right). **d**, Western blot detection of known nuclear RBPs in samples from (**c**). SRSF1 and hnRNPC are known nuclear RBPs, GRSF1 is a mitochondrial RBP, and ETS2 is a nuclear protein that does not bind RNA (non-RBP).

To test and optimize the APEX-PS protocol, we first focused on nuclear RBPs, which constitute a well-studied class of subcellular RBPs that have been characterized by fractionation-based methods, such as serIC (Serial RNA interactome capture)^32^ and RBR-ID (RNA-binding region identification)^33^. We prepared HEK293T cells stably expressing nucleus-targeted APEX-NLS, and performed live-cell labeling with biotin-phenol for 1 minute. We then treated the cells with FA for 10 minutes, lysed, and performed PS followed by streptavidin bead enrichment. **Fig. 1c** shows that many biotinylated proteins were recovered by this procedure but not in negative controls omitting H_2_O_2_ (which suppresses APEX-catalyzed biotinylation) or FA (which prevents RNA-protein crosslinking and hence recruitment of proteins to the interphase).

To probe the identities of nuclear APEX-PS enriched proteins, we performed Western blotting of enriched lysates (**Supplementary Fig. 1a**) for known nuclear RBPs as well as off-target protein markers (**Fig. 1d**). We detected the nuclear RBPs hnRNPC (heterogeneous nuclear ribonucleoproteins C1/C2) and SRSF1 (serine/arginine-rich splicing factor 1), while the mitochondrial RBP GRSF1 (G-rich sequence factor 1) and nuclear non-RBP ETS2 (C-ets-2) were not detected. Notably, GRSF1 was enriched after the PS step, because it is an RBP, but not after streptavidin bead enrichment, because it is not a nuclear protein and thus not biotinylated by APEX-NLS.

To explore the generality of APEX-PS, we repeated the protocol for four additional subcellular compartments: the cytosol, ER membrane (facing the cytosol), outer mitochondrial membrane (facing the cytosol), and nucleolus (**Supplementary Fig. 1b**). All of these gave H_2_O_2_- and FA-dependent signals in streptavidin blots of whole cell lysates, with distinct banding patterns or “fingerprints” unique to their compartment (**Supplementary Fig. 1c-f**). We also tested the APEX-PS workflow with UV crosslinking instead of FA (**Supplementary Fig. 1g**). Despite reduced protein recovery due to lower UV crosslinking efficiency (**Supplementary Fig. 1h**), we could again specifically enrich nuclear RBPs over non-nuclear RBPs and nuclear non-RBPs using APEX-NLS-expressing cells (**Supplementary Fig. 1i**). Finally, we tested proximity labeling with TurboID^17^ instead of APEX2 (**Supplementary Fig. 1j**; TurboID-PS). Streptavidin blotting of enriched lysates from TurboID-NES-expressing HEK293T cells showed biotin- and FA-dependent recovery of endogenous RBPs. Using TurboID instead of APEX for compartment-specific RBP discovery may be preferable in *in vivo* systems due to the simplicity of biotin addition and avoidance of H_2_O_2_ toxicity.

### Proteomic profiling of nuclear RBPs by APEX-PS

Having validated APEX-PS in the nucleus by Western blot, we proceeded to a multiplexed, TMT-based proteomics experiment that included both nucleus-targeted APEX2-NLS and nucleolus-targeted APEX2-NIK3x (the latter discussed below). The experimental design (**Fig. 2a**) consisted of three biological replicates of each construct expressed in HEK293T cells, alongside three negative controls with APEX2, H_2_O_2_, or FA omitted. Imaging showed correct localization of both APEX2-NLS (V5 staining) and APEX-biotinylated proteins (neutravidin staining) (**Fig. 2b**). After phase separation, streptavidin enrichment, and on-bead tryptic digestion to peptides, each sample was chemically tagged with a unique TMT label. The samples were then pooled and analyzed by liquid chromatography-tandem mass spectrometry analysis (LC-MS/MS).

**Fig. 2.**
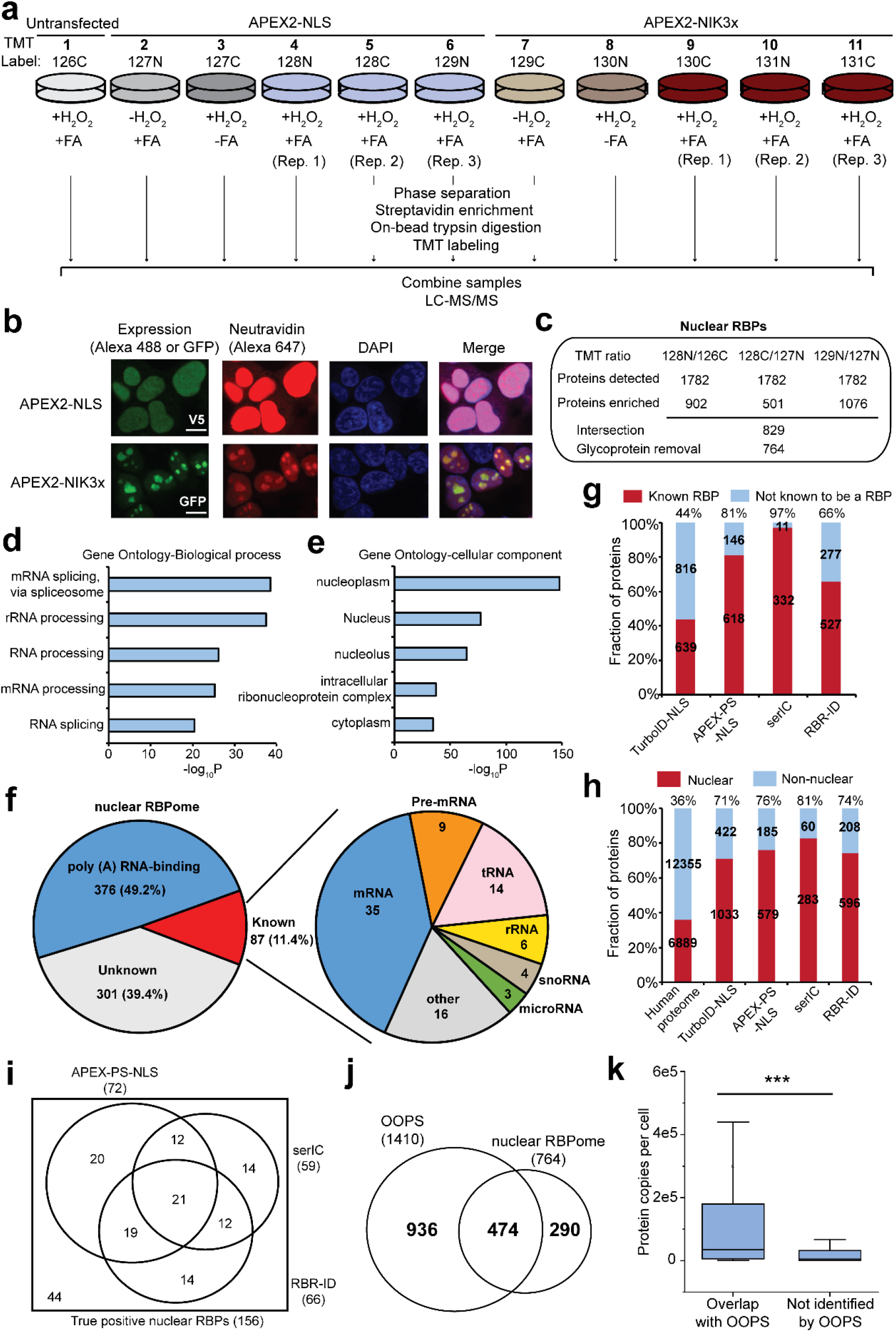
Proteomic profiling of nuclear RBPs by APEX-PS. **a**, Experimental design and labeling conditions for TMT-based proteomics. HEK293T cells stably expressing nuclear APEX2-NLS (samples 2-6) or nucleolar APEX2-NIK3x (samples 7-11) were subjected to proximity biotinylation and FA crosslinking. The control samples omitted enzyme (sample 1), H_2_O_2_ (sample 2 and 7), or FA (3 and 8). **b**, Confocal fluorescence imaging of APEX2 localization (V5 or GFP) and biotinylation activity (neutravidin-Alexa 647) in the nucleus (top) and nucleolus (bottom). Live cell biotinylation was performed for 1 minute before fixation. DAPI stains nuclei. Scale bars, 10 μm. **c**, Numbers of proteins remaining after each step of filtering the mass spectrometry data. The final nuclear RBPome obtained by APEX-PS has 764 proteins (Supplementary Dataset 2). **d**, GO biological process analysis of nuclear RBPs identified by APEX-PS. **e**, GO cellular component analysis of nuclear RBPs identified by APEX-PS. **f**, Subclassification of RBPs in nuclear APEX-PS dataset. Many RBPs we enriched bind to poly(A) ^6, 7, 12, 32, ^34, 35^^. Of the remainder, 87 have been experimentally shown to bind to the 7 RNA classes at right. Details in Supplementary Dataset 2. **g**, RBP specificity of nuclear datasets. For comparison, the nuclear proteome identified by TurboID-NLS^17^ and nuclear RBPs identified by fractionation^32, 33^ were analyzed in the same manner. Details in Supplementary Dataset 2. **h**, Nuclear specificity of nuclear RBP datasets identified by APEX-PS, compared to previous TurboID-NLS dataset and nuclear RBPs identified by fractionation (RBR-ID and serIC). **i**, Using a list of 156 true-positive nuclear RBPs, the coverage of APEX-PS was compared to fractionation-based methods. The nuclear RBPome consists of 764 proteins enriched in 2 or more of the 3 experimental replicates. **j**, Overlap of APEX-PS datasets with global RBP dataset obtained by OOPS^12^. **k**, Comparison of protein abundance for RBPs identified by both methods to those identified by APEX-PS only.

A total of 1782 proteins were detected (each with two or more unique peptides) (**Fig. 2c and Supplementary Dataset 1**). We observed excellent correlation across the three biological replicates (**Supplementary Fig. 2a**, r^2^ = 0.91). To filter the data and arrive at our final nuclear RBPomes, we first paired each replicate with a negative control to calculate TMT ratios. We then used a true positive list of known nuclear RBPs and a false positive list of non-nuclear proteins (which should not be biotinylated by APEX2-NLS) to generate receiver operating characteristic (ROC) curves for each replicate (**Supplementary Fig. 2b**). We applied the TMT ratio cutoff that maximized retention of true positives and minimized retention of false positives for each dataset (**Supplementary Fig. 2c**). From the three resulting protein lists, we included in our “nuclear RBPome” the 829 proteins that were enriched in two or more replicates (when we used the more stringent criterion of enrichment in all three replicates, the specificity did not improve while the sensitivity decreased). Because phase separation is known to enrich glycosylated proteins due to the hydrophilicity of glycans ^12^, we manually removed 65 secretory pathway glycoprotein contaminants found in our nuclear RBPome, to give a final proteome size of 764 proteins (**Supplementary Dataset 2**).

### Sensitivity and specificity of nuclear RBPome generated by APEX-PS

Gene Ontology (GO) analysis of our nuclear RBPome showed enrichment of RNA-related terms including mRNA splicing and processing (**Fig. 2d**). GO cellular component (GOCC) analysis showed significant enrichment of nucleus-related terms, as expected (**Fig. 2e**). We crossed our dataset with previous polyadenylated RBP datasets^6, 7, 12, 32, 34, 35^ and determined that 48% of our RBPs bind to polyadenylated RNAs while the remainder bind to tRNA, rRNA, snoRNA, etc. or have unknown RNA binding partners (**Fig. 2f**).

To benchmark our methodology against prior approaches, we considered the nuclear RBP datasets obtained by serIC ^32^ and RBR-ID ^33^. serIC combines nuclear fractionation with repeated oligo(dT) RBP purification; 343 nuclear RBPs were enriched by this method from 1 × 10^9^ cells (20 times more material than used for APEX-PS). RBR-ID is based on depletion of parent peptide signatures in MS due to RNA-protein crosslinking. 804 nuclear RBPs were identified from purified mouse embryonic stem cell nuclei by this approach.

Specificity was determined by calculating the fraction of each dataset with prior RNA binding (**Fig. 2g**) or nuclear (**Fig. 2h**) annotation. Sensitivity was calculated by first manually assembling a list of 156 well-established nuclear RBPs involved in fundamental processes such as ribosome biogenesis, splicing and transcription (**Supplementary Dataset 2**). We then asked what fraction of our “true positive” list was detected by each method. We found that serIC was extremely specific (97% RBP specificity) but less sensitive (38%), missing RBPs that bind to non-polyadenylated RNAs (e.g., tRNA ligases) for example (**Supplementary Fig. 2d**). RBR-ID was more sensitive than SerIC (42%) but much less specific (66% RBP specificity). Our APEX-PS dataset, quantified in the same manner, showed reasonable specificity (81% RBP specificity) and the highest sensitivity of the three methods (46%). The true specificity of APEX-PS may be higher, as many of the “orphans” we identified (proteins lacking prior RBP annotation) may represent novel nuclear RBPs. For instance, we enriched many orphan proteins related to transcription and DNA repair, which may represent new RBPs involved in genome regulation (**Supplementary Fig. 2e**). The Venn diagram comparing the “true positive” proteins identified by the three methods is shown in **Fig. 2i** and APEX-PS provided the largest dataset (72 proteins), starting with the smallest amount of cellular material.

We also compared nuclear APEX-PS to the previous global OOPS dataset in HEK293T cells ^12^ (**Fig. 2j**). Interestingly, APEX-PS identified 290 nuclear RBPs missed by OOPS; these tend to be lower-abundance proteins^36^ (**Fig. 2k**). We surmise that by focusing on a single subcellular compartment, APEX-PS can probe more deeply than global profiling methods and identify lower-abundance RBPs.

### Identification of nucleolar RBPs by APEX-PS

Having validated APEX-PS in the nucleus, we turned our attention to the nucleolus, a membraneless subcompartment that is inaccessible to fractionation-based approaches for RBP discovery such as SerIC and RBR-ID. We performed three biological replicates of APEX-PS in HEK293T cells stably expressing APEX2 targeted to the nucleolus (**Fig. 2a, b**). To maximize spatial specificity, we performed ratiometric analysis^37^, referencing the whole-nucleus APEX2-NLS samples (**Fig. 3a, b and Supplementary Fig. 3a**). After intersection of three independent datasets and removal of three potential glycoprotein contaminants, we arrived at a nucleolar RBPome of 252 proteins (**Table S3**).

**Fig. 3.**
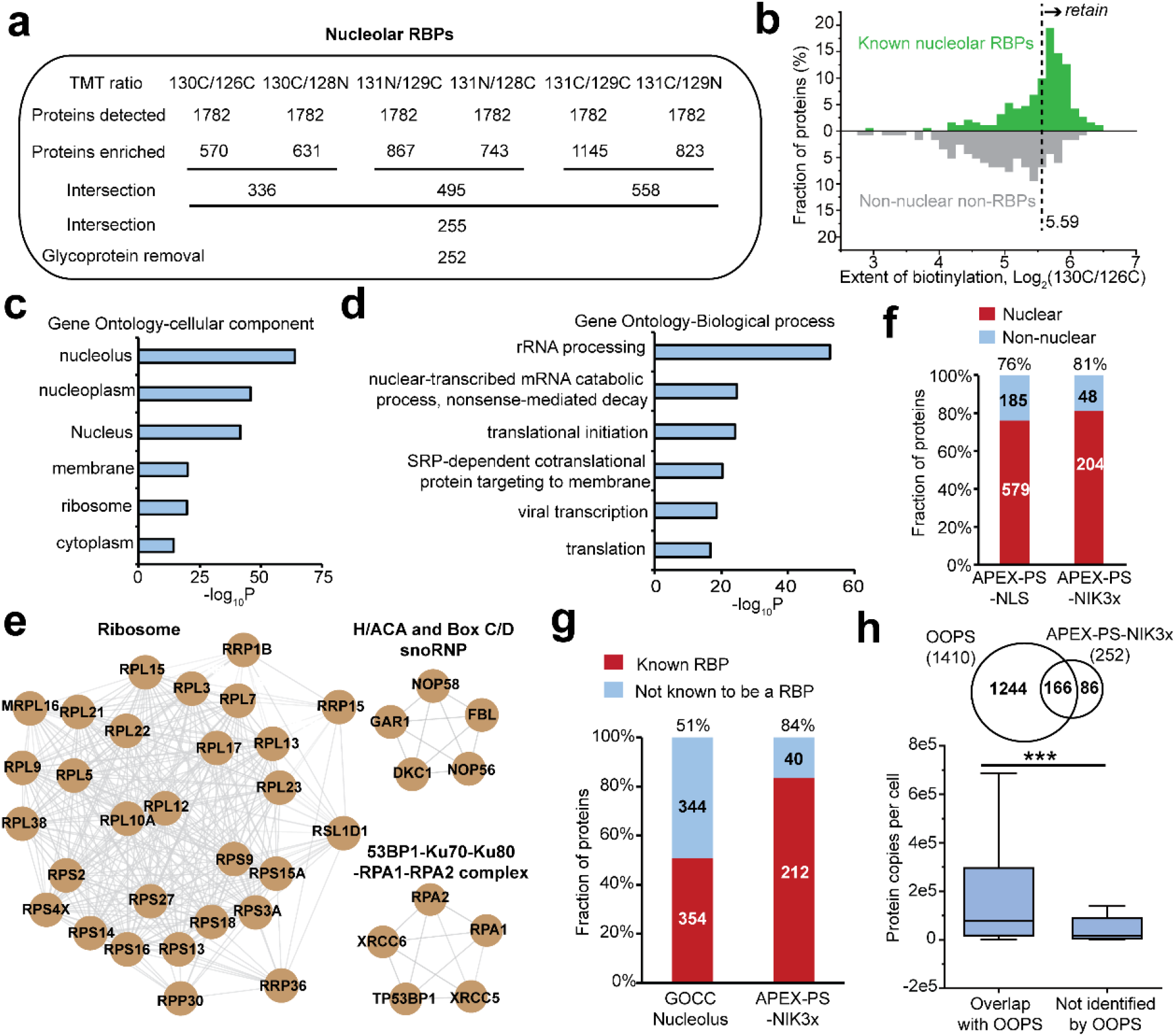
Proteomic profiling of nucleolar RBPs by APEX-PS. **a**, Numbers of proteins remaining after each step of filtering the mass spectrometric data. The final nucleolar RBPome has 252 proteins (Supplementary Dataset 3). **b**, Sample histogram showing how the cutoff for 130C/126C ratio was applied. **c**, GO cellular component analysis of nucleolar RBPs identified by APEX-PS. **d**, GO biological process analysis of nucleolar RBPs identified by APEX-PS. **e**, Molecular complexes enriched in nucleolar RBPome. Gray lines indicate protein-protein interactions annotated by the STRING database. **f**, Nuclear specificity of nucleolar RBPome. **g**, RBP specificity of nucleolar dataset. For comparison, the 698 proteins with nucleolus annotation in GOCC were analyzed in the same manner. See Supplementary Dataset 3 for details. **h**, Overlap with global RBP dataset obtained by OOPS^12^. Bottom: Comparison of protein abundance for RBPs identified by both methods to those identified by APEX-PS only.

Gene ontology analysis showed enrichment of expected cell compartment (**Fig. 3c**) and biological process (**Fig. 3d**) terms, including rRNA processing and translation initiation, consistent with the understanding that the nucleolus is the primary site for ribosome biogenesis. In addition to enriching 29 ribosomal protein subunits, we also detected several H/ACA and Box C/D small nucleolar ribonucleoprotein complexes (snoRNP), which catalyze the pseudouridylation and 2’-O-ribose methylation of rRNA respectively^38^ (**Fig. 3e**). 81% and 84% of our nucleolar proteins have prior nuclear and RNA binding annotation, respectively (**Fig. 3f, g**). Several of the unannotated orphans have connections to cell division and ubiquitination **(Supplementary Fig. 3b**). Comparing again to global OOPS^12^, we found that 86 of our nucleolar RBPs were missed by OOPS, and that these are generally lower-abundance proteins (**Fig. 3h**). Similarly, 92 nucleolar RBPs were missed by poly (A)-dependent methods^6, 7, 12, 32, 34, 35^, perhaps because they bind to non-coding RNAs (**Supplementary Fig. 3c**).

### Analysis of RNA-binding domains reveals RBPs with high RNA binding affinity

RBPs bind RNA via modular RNA binding domains (RBDs) that have been classified into 11 classical and 15 non-classical sub-types^7, 39^. In our nuclear RBPome, more of the proteins with prior RNA binding annotation have classical than non-classical RBDs (**Fig. 4a, b**). By contrast, among our RBP orphans, far more have non-classical RNA binding domains than classical. A similar trend was observed for our nucleolar RBPome (**Supplementary Fig. 4a, b**). This suggests that RBPs with non-classical RBDs may be less explored than RBPs with classical RBDs, and that APEX-PS may facilitate discovery of the former. Notably, consistent with previous RBP studies, a large percentage of RBPs that we mapped do not have characterized RBDs, suggesting that they may bind RNA through novel mechanisms.

**Fig. 4.**
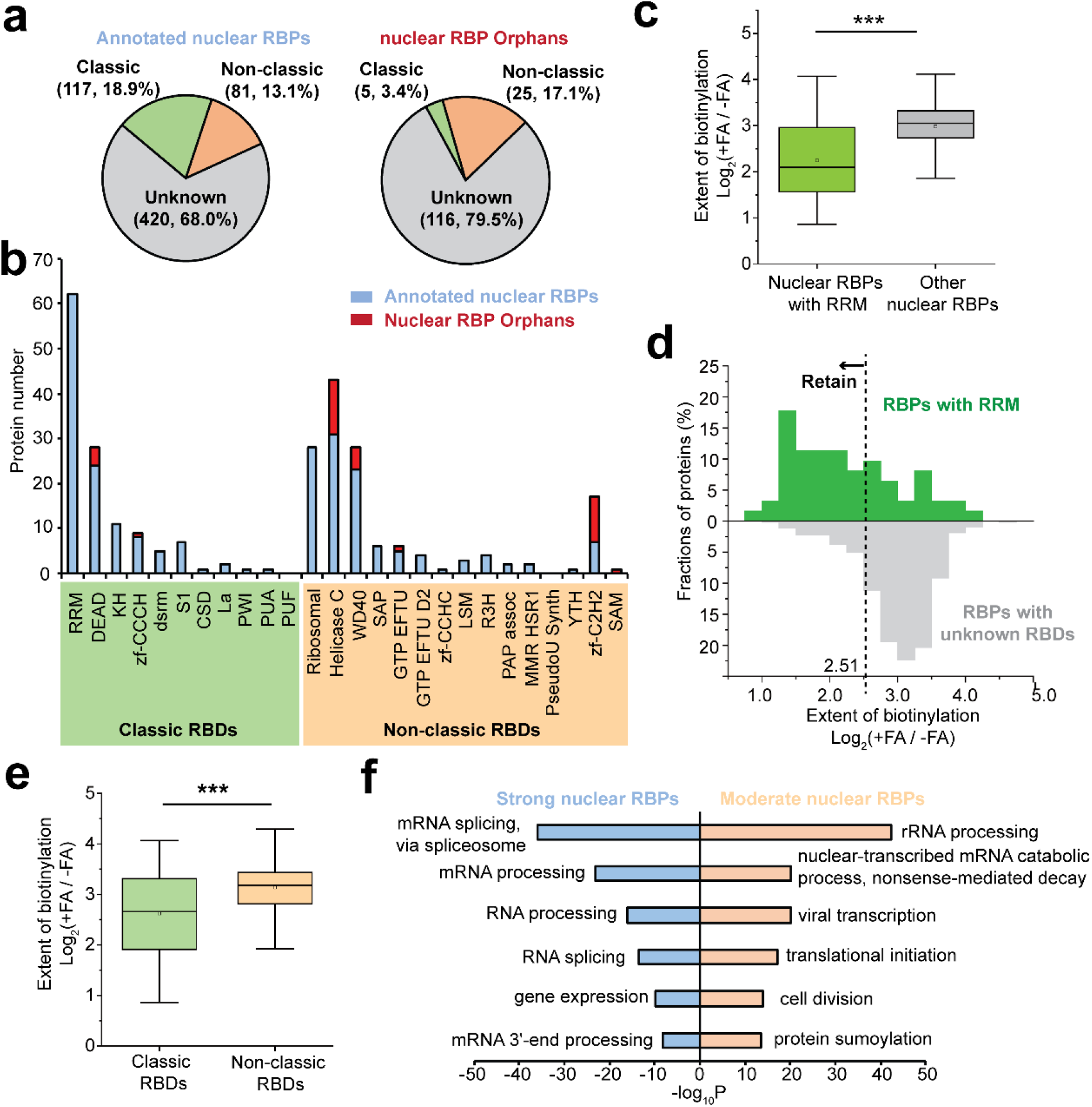
RNA binding domain (RBD) analysis. **a**, Percentages of APEX-PS detected nuclear RBPs with classical versus non-classical RNA binding domains (RBDs). **b**, Number of proteins with each type of classical or non-classical RBD. The RBDs of each RBP are listed in Supplementary Dataset 2. **c**, Comparison of formaldehyde-dependent APEX-PS enrichment for RRM-containing RBPs (green) versus unknown (grey)-RBD-containing proteins in the nuclear APEX-PS dataset. **d**, Histogram showing distribution of RRM (green) versus unknown (grey)-RBD-containing proteins in the nuclear APEX-PS dataset by formaldehyde-dependent enrichment. **e**, Comparison of formaldehyde-dependent APEX-PS enrichment for classical (green) versus non-classical (yellow)-RBD-containing proteins in the primary nuclear APEX-PS dataset. **f**, GO biological process analysis of strong (low +FA/-FA ratio) and moderate (medium +FA/-FA ratio) nuclear RBPs.

In the nuclear and nucleolar proteomic experiment depicted in **Fig. 2a**, we included negative controls with the FA crosslinking step omitted. However, our mass spectrometric data showed that these conditions still enriched some RBPs. We wondered whether these RBPs might represent tighter-binding RBPs that remain bound to their RNA partners through cell lysis and phase separation, even in the absence of FA crosslinking. To assess this, we performed analysis of detected proteins based on +FA/-FA enrichment ratio. We found that RBPs containing the RNA recognition motif (RRM), which interacts tightly (nanomolar affinity) with RNA binding partners via multiple sequential stacking interactions^2, 40^, generally displayed low +FA/-FA enrichment ratios, consistent with the idea that high-affinity RBPs can be enriched by phase separation even in the absence of chemical crosslinking (**Fig. 4c, d**). We also observed that the distribution of RBPs with classical RBDs were shifted towards lower +FA/-FA ratio than RBPs with non-classical RBPs; this suggests that classical RBDs may tend to bind RNA with higher affinity (**Fig. 4e and Supplementary Fig. 4c**). Gene ontology analysis of the 148 RBPs with lower +FA/-FA ratio (higher RNA affinity) and 616 RBPs with higher +FA/-FA ratio (lower RNA affinity) enriched distinct biological process terms, including spliceosome for the former and rRNA processing for the latter. (**Fig. 4f)**

### Discovery of RBPs at the outer mitochondrial membrane by APEX-PS

The landscape of RBPs at the mammalian outer mitochondrial membrane (OMM) has not previously been explored but could hold clues to the mechanisms of mitochondrial protein translation. Of the >1200 proteins assigned to the mitochondrion, only 13 are encoded by the mitochondrial genome. The remainder are encoded by the nuclear genome, and their protein products must be imported into the mitochondrion after translation in the cytosol. Since the detection of ribosomes at the OMM by electron microscopy^41^, several studies, including ours^24, 42^, have detected mitochondrial mRNAs at the OMM, suggesting that these may be translated “locally” to facilitate co-translational or post-translational import of their protein products in the mitochondrion. For instance, our APEX-seq study detected 1902 OMM-localized mRNAs in HEK cells (Fazal et al., 2019), while proximity-specific ribosome profiling showed active translation of 551 mRNAs at the yeast OMM ^24, 42^. These observations raise the question of how mitochondrial mRNAs are recruited to the OMM for local translation – are specific OMM-localized RBPs involved?

To identify OMM-localized RBPs, HEK293T cells stably expressing APEX2-OMM were subjected to proximity biotinylation, FA crosslinking, and phase separation in two biological replicates, while cytosolic APEX2-NES was used as a spatial reference control in a TMT 11-plex experiment (**Fig. 5a, Supplementary Fig. 5a, b and Supplementary Dataset 4**). We also included two replicates of cells treated with puromycin (PUR), which inhibits protein translation and disassembles polysomes^43^, in order to detect OMM-localized RBPs that bind to RNA in a translation and ribosome-independent fashion (**Fig. 5b**).

**Fig. 5.**
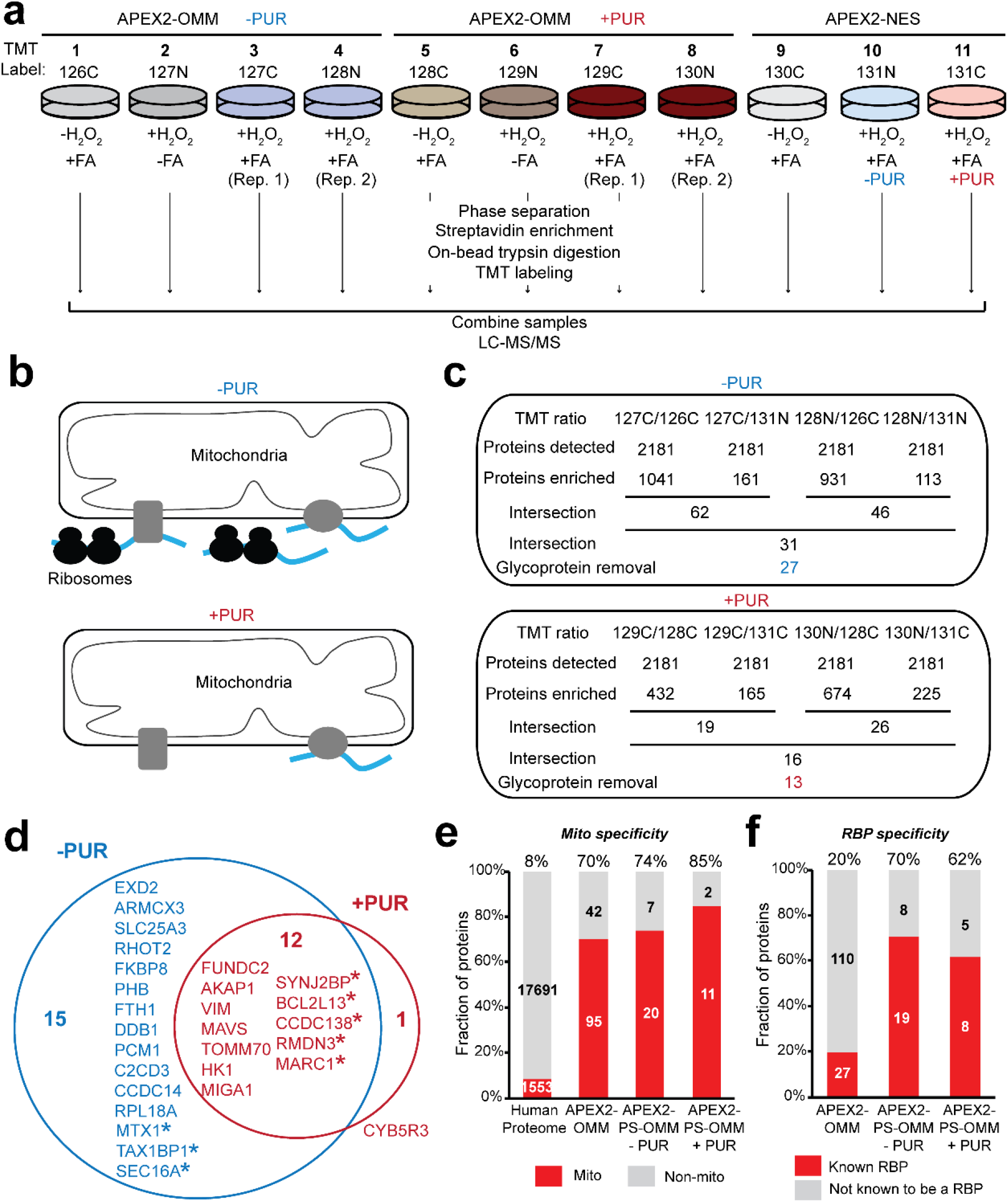
Proteomic profiling of RBPs localized to the outer mitochondrial membrane by APEX-PS. **a**, Experimental design and labeling conditions for TMT-based proteomics. HEK cells stably expressing APEX2-OMM were subjected to proximity biotinylation, FA crosslinking, and enrichment according to Figure 1a-b. The control samples omitted H_2_O_2_ (samples 1 and 5), or FA (samples 2 and 6). Cytosolic APEX2-NES was included as a reference for ratiometric analysis (samples 9-11). **b**, Schematic showing expected changes at the OMM upon treatment with puromycin (PUR), which inhibits protein translation and disrupts polysomes. Blue lines, RNA. Grey shapes, potential RBPs. **c**, Numbers of proteins remaining after each step of filtering the mass spectrometric data. The final nuclear –PUR and +PUR OMM RBP proteomes obtained by APEX-PS have 27 and 13 proteins, respectively. **d**, Overlap of OMM-localized RBPs under basal and +PUR conditions. RBP orphans (proteins with no previous RNA binding annotation in literature) are starred. **e**, Mitochondrial specificity of OMM RBPs identified by APEX-PS. For comparison, same analysis performed on APEX2 OMM proteome^45^ and entire human proteome. Details in Supplementary Dataset 5. (**F**) RBP specificity of APEX-PS OMM datasets. Details in Supplementary Dataset 5.

A total of 27 and 13 OMM-localized RBPs were identified under basal and PUR-treated conditions, respectively (**Fig. 5c and Supplementary Dataset 5**). The +PUR dataset is almost entirely a subset of the basal (-PUR) dataset (**Fig. 5d**). Most proteins (75%) in our OMM RBPome have prior mitochondrial annotation, and 60% have prior RNA binding annotation (**Fig. 5e, f**). Interestingly, several proteins (8 out of 27) also have literature connections to mitochondria-ER contact sites, which previous studies have linked to protein translation^44-46^ (**Supplementary Fig. 5c**).

We selected two proteins for follow-up validation. SYNJ2BP was enriched under both basal and +PUR conditions and is a tail-anchored OMM protein with a cytosol-facing PDZ domain. We previously showed that the complex of SYNJ2BP and its ER binding partner, RRBP1, functions as a mitochondria-rough ER tether^45^. SYNJ2BP has not previously been shown to bind RNA but, interestingly, the PDZ domain has been suggested to have RNA-binding activity^34^. The second protein we selected, EXD2, was enriched at the OMM under basal conditions only. EXD2 is an OMM-localized 3’-5’ exonuclease that acts on single-stranded RNA^47^ and has been shown to influence intra-mitochondrial translation^48^, perhaps by acting on cytosolic mRNAs encoding mitochondrial ribosome subunits. EXD2, like SYNJ2BP, has also been detected at mitochondria-ER contact sites^46, 49^. To validate both these proteins as bona fide RBPs, we first repeated APEX-PS but used UV crosslinking (shorter range, but more specific) instead of FA-based crosslinking. The Western blot in **Fig. 6a** shows enrichment of both these proteins by APEX-PS (UV) at the OMM. Second, we used an orthogonal RBP enrichment method consisting of RNA metabolic labeling followed by click chemistry and pull down, which again enriched both proteins (**Fig. 6b**).

**Fig. 6.**
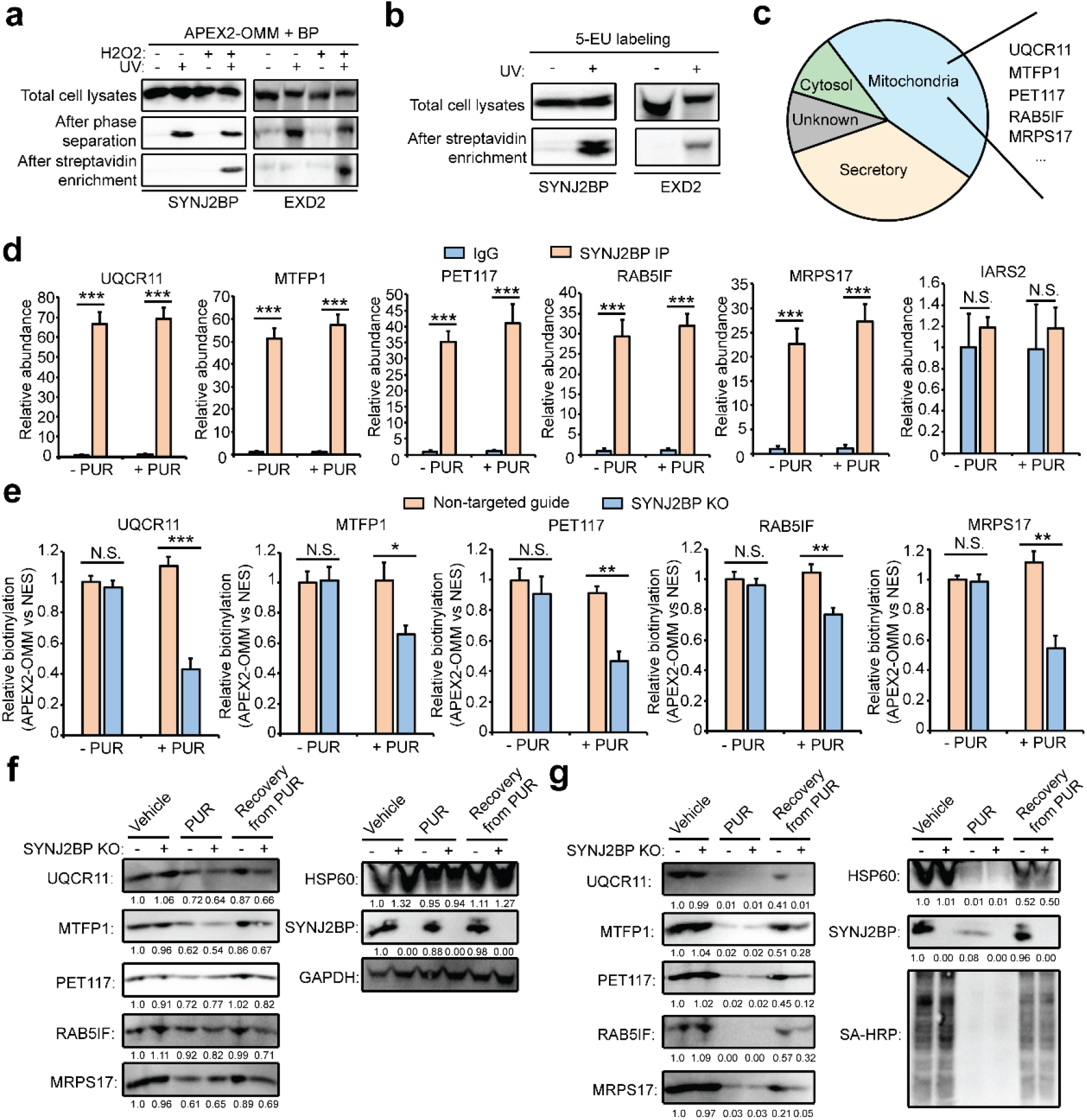
SYNJ2BP binds and promotes the translation of a subset of OMM-localized mRNAs. **a**, Validation of SYNJ2BP and EXD2 as novel OMM-localized RBPs by UV crosslinking APEX-PS. **b**, Validation of the RNA binding of SYNJ2BP and EXD2 by metabolic labeling of RNA by 5-EU and UV crosslinking. **c**, The subcellular locations of proteins encoded by top20 SYNJ2BP mRNA targets. **d**, Validation of mitochondria-related SYNJ2BP clients in the absence or presence of protein translation inhibitor PUR by CLIP and qRT-PCR. IARS2 is an OMM-localized mRNA not identified to bind with SYNJ2BP. **e**, Evaluation of OMM localization of SYNJ2BP clients in SYNJ2BP knock-out cells by APEX-mediated proximity labeling of RNA. The OMM localization was determined by comparing APEX2-OMM with APEX2-NES labeling. **f**, Evaluation of the impact of SYNJ2BP knock-out on the protein level of SYNJ2BP-regulated clients under different cellular states. **g**, Evaluation of the impact of SYNJ2BP knock-out on protein synthesis of SYNJ2BP-regulated clients under different cellular states. The newly synthesized proteins were labeled by AHA and captured by click-based enrichment.

### Analysis of the mRNA clients of the OMM-localized RBP SYNJ2BP

We were intrigued by SYNJ2BP and especially its persistence as an RBP after puromycin treatment, suggesting that it binds to RNA in a ribosome- and translation-independent manner. To further probe the function of SYNJ2BP, we performed RNA immunoprecipitation and sequencing (RIP-seq) to identify the mRNA targets of SYNJ2BP. More than 100 mRNAs were highly enriched, most of them encoding mitochondrial and secretory proteins (**Fig. 6c and Supplementary Dataset 6**). We selected 11 mRNAs with distinct enrichment for validation by cross-linking immunoprecipitation (CLIP) and RT-qPCR. **Fig. 6d and Supplementary Fig. 6a** shows that all 11 mRNAs were enriched by CLIP-based pull-down of endogenous SYNJ2BP, while negative controls including IARS2, an OMM-localized mRNA ^24^ that was not detected by SYNJ2BP RIP-seq, were not enriched. Following PUR treatment, all 11 mRNA clients were enriched to a similar extent, suggesting that SYNJBP binds to these mRNAs regardless of ribosome activity. Imaging confirmed that SYNJ2BP remains OMM-localized after PUR treatment (**Supplementary Fig. 6b**).

To determine if these 11 mRNAs are localized to the OMM solely through the action of SYNJ2BP or if other binding interactions also play a role, we used OMM-localized APEX to directly biotinylate the RNAs at the OMM in both wild-type and SYNJ2BP knock-out HEK293T cells (**Supplementary Fig. 6c**). We found that 5 of the 11 mRNAs were significantly reduced at the OMM when SYJN2JBP was absent (**Fig. 6e**). This effect was specific to the PUR-treated condition only. Our observations suggest that binding to SYNJ2BP is a major mechanism for retention of a specific subset of mRNAs at the OMM following PUR treatment, but other mechanisms exist for recruiting these mRNAs to the OMM under basal conditions – for instance, interactions between the TOM mitochondrial protein import machinery and the ribosome-mRNA-nascent protein chain complex that displays an N-terminal mitochondrial targeting sequence^50^. The five SYNJ2BP-dependent mRNAs consist of three oxidative phosphorylation (OXPHOS)-related proteins (UQCR11, PET117 and RAB5IF), a mitochondrial ribosome component (MRPS17) and a key mitochondrial fission factor (MTFP1).

### SYNJ2BP retains specific mitochondrial mRNAs at the OMM to facilitate restoration of mitochondrial function after stress

Due to the functional importance of these genes, we hypothesized that SYNJ2BP’s role may be to retain them at the OMM, even through periods of stress, in order to facilitate their rapid local translation for restoration of mitochondrial function and/or biogenesis. To test this hypothesis, we examined the effect of SYNJ2BP KO on the levels of proteins encoded by these genes, under basal conditions, after PUR treatment, and after 12 hours of recovery from PUR treatment. No change in protein levels were observed under basal and PUR conditions, but all five proteins were reduced in abundance upon SYNJ2BP KO 12 hours after recovery from PUR (**Fig. 6f**). Because long protein half-lives may obscure the full effect of SYNJ2BP on protein translation, we repeated the assay but used azidohomoalanine (AHA) (followed by Click reaction with alkyne-biotin and streptavidin-based enrichment ^51^) to selectively detect newly synthesized proteins. **Fig. 6g** shows that SYNJ2BP KO clearly impaired the translation of all five proteins during the PUR recovery phase. In a control experiment, SYNJ2BP had no impact on the total protein level or new protein level of HSP60, a mitochondrial protein whose mRNA is not a client of SYNJ2BP.

To investigate whether SYNJ2BP’s role in mitochondrial protein translation has an effect on overall mitochondrial function, we first performed measurements of OXPHOS, specifically the activities of Complex III and Complex IV. One of SYNJ2BP’s validated mRNA clients, UQCR11, encodes a subunit of Complex III^52^, the component of the electron transport chain that catalyzes electron transfer from ubiquinol to cytochrome c. **Fig. 7a** shows that PUR treatment, unsurprisingly, impairs the activity of Complex III. During the recovery from PUR stress, the Complex III activity of wild-type cells returns to previous levels, while the Complex III activity of SYNJ2BP KO cells fails to recover. We also measured the activity of Complex IV, whose assembly requires PET117, another validated SYNJ2BP client. Previous work has suggested that PET117 is essential for OXPHOS and PET117 mutation leads to neurodevelopmental regression due to a deficiency in Complex IV assembly^53, 54^. **Fig. 7b** shows PUR-induced decrease in Complex IV activity, and failure to return to previous levels of activity during stress recovery, only in SYNJ2BP KO cells but not in wild-type cells.

**Fig. 7.**
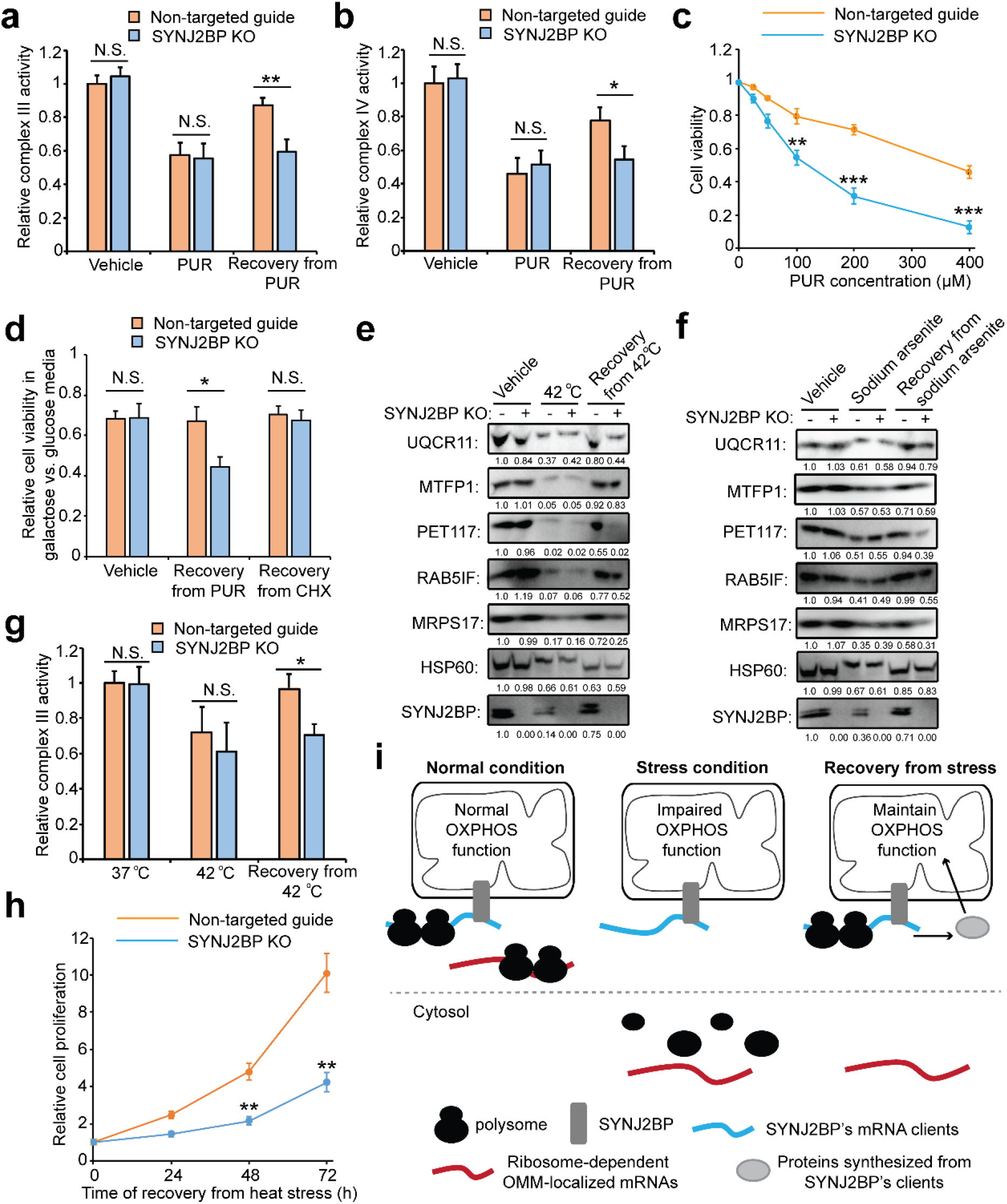
SYNJ2BP promotes the cellular recovery from stresses. **a-b**, Evaluation of complex III (**a**) and IV (**b**) activity in SYNJ2BP knock-out cells. **c**, Evaluation of cell viability pre-treated with protein translation inhibitor PUR. **d**, Preferential utilization of galactose versus glucose as a carbon source in SYNJ2BP knock-out cells. **e-f**, Evaluation of protein synthesis of SYNJ2BP clients in SYNJ2BP knock-out cells under heat (**e**) and sodium arsenite (**f**) stresses. **g**, Evaluation of complex III activity in SYNJ2BP knock-out cells under heat stress. **h**, Evaluation of cell proliferation in SYNJ2BP knock-out cells during the recovery from heat stresses. **i**, The working model of SYNJ2BP in mediating the local translation of OMM-localized mRNAs during the recovery from stresses.

To interrogate the effect of SYNJ2BP on mitochondrial health overall, we performed measurements of cell proliferation, which depends on mitochondrial activity, especially when cells are cultured in galactose. Cells grown in glucose derive ATP from both aerobic glycolysis and mitochondrial glutamine oxidation, whereas cells growth in galactose derive a larger fraction of their ATP from mitochondrial respiration^55^. We found that in glucose media, WT and SYNJ2BP KO cells grew identically under basal conditions, but SYNJ2BP KO cells grew more slowly than wild-type after PUR treatment (**Fig. 7c and Supplementary Fig. 7a**). This effect was not observed using cycloheximide (CHX), which also inhibits protein translation but keeps polysomes intact (**Supplementary Fig. 7b**). In the presence of CHX, the OMM-localized mRNAs could be retained by polysomes and thus cells survived in a SYNJ2BP-independent manner. When the cells were grown in galactose to increase reliance on mitochondrial respiration, the effect of SYNJ2BP KO on cell growth following PUR treatment became more pronounced (**Fig. 7d**). We also measured ATP levels directly and found that SYNJ2BP KO decreased ATP in galactose-cultured cells after PUR treatment, but not under basal or CHX-treated conditions (**Supplementary Fig. 7c**). Collectively, our results suggest that SYNJ2BP retains specific mitochondrial transcripts at the OMM following PUR-specific translation inhibition stress, and that this serves to improve the recovery of OXPHOS, mitochondrial ATP production, and cell growth from such stress.

Apart from PUR treatment, other types of stresses, such as heat and oxidative stress, can also suppress translation and dissociate mRNAs from polysomes^56^. The released mRNAs interact extensively with RBPs, inducing the formation of molecular condensates such as stress granules^57^. To test if SYNJ2BP also plays a role in the recovery of mitochondrial function from other types of translation stress, we first used heat or sodium arsenite to stress cells, and then checked for SYNJ2BP-dependent retention of five mRNA clients by OMM-APEX-seq. **Supplementary Fig. 7d** shows that all five mRNAs associate with the mitochondrion under heat and sodium arsenite stress in wide-type cells, but largely disappear from the OMM in SYNJ2BP knock-out cells. Moreover, loss of SYNJ2BP also inhibits the translation of these mRNAs under both stresses, which further leads to the inhibition of Complexes III and IV (**Fig. 7e-g and Supplementary Fig. 7e-g)**. In a proliferation assay, SYNJ2BP knock-out impaired cell growth during recovery from heat and sodium arsenite stresses (**Fig. 7h and Supplementary Fig. S7h)**. Taken together, SYNJ2BP may represent a novel mechanism for the specific regulation of local translation at the OMM in response to cellular stresses (**Fig. 7i)**.

## Discussion

RNA binding proteins (RBPs) constitute an important subclass of the human proteome, with essential roles in transcription, translation, chromatin organization, innate immunity, metabolite sensing, and stress response. Here we showed how proximity labeling with promiscuous enzymes can be used for unbiased discovery of RBPs in specific subcellular compartments. By combining APEX-mediated proximity biotinylation with organic-aqueous phase separation to enrich crosslinked protein-RNA complexes, we were able to map nuclear RBPs in human cells with high specificity and sensitivity compared to previous fractionation-based methods. We showed that APEX-PS can also be applied for RBP discovery in unpurifiable subcompartments, such as the nucleolus and outer mitochondrial membrane (OMM), which are inaccessible to biochemical fractionation. Taking advantage of the ∼11 minute temporal resolution of APEX-PS, we compared the RBPome of the OMM before and after translation inhibition stress. By doing so, we identified a novel RBP that protects a specific subset of mitochondrial mRNAs from relocalization away from the mitochondrion during stress. This retention mechanism appears to facilitate local protein synthesis from these transcripts during stress recovery, giving rise to rapid restoration of OXPHOS, ATP synthesis, and overall mitochondrial function.

Many innovative strategies for large-scale discovery of RBPs have been reported in recent years. UV/formaldehyde crosslinking followed by oligo(dT) bead capture can enrich RBPs that bind to polyadenylated RNAs^6-9^. Metabolic labeling of RNA with alkyne analogs used in CARIC^10^and RICK^11^ identifies polyadenylation-independent RBPs. Several groups have reported organic-aqueous phase separation for RBP enrichment independent of RNA class^12-14^. Notably, phase separation approaches feature ∼100-fold greater sensitivity than oligo(dT) pulldown^12^. While these methods have uncovered thousands of newly annotated RBPs, the vast majority of them have unknown or partially known function. As a first step in elucidating their biology, the spatial assignment of RBPs to specific subcellular locales, under specific cellular states, could help to shed light on their functional roles.

To recover spatial information, previous studies have crossed RBP datasets with protein localization datasets^58^, even though such datasets are often acquired in different cell types under different conditions. Moreover, many RBPs reside in multiple subcellular locations but bind to RNA in only one of these locations; such information is lost by dataset crossing. Another, more direct, approach to spatial assignment of RBPs is to combine RBP profiling methods with subcellular fractionation. This is effective for organelles that can be enriched by biochemical fractionation, such as the nucleus^59^, but inapplicable to the many subcellular regions that are impossible to purify, such as stress granules, processing bodies, and the nucleolus and OMM that we map here with APEX-PS.

Proximity labeling (PL) with enzymes such as APEX, TurboID, and BioID has been widely applied and is straightforward to implement^19^. Here, the addition of two simple steps - crosslinking and phase separation - results in a dataset that has *functional* annotation in addition to the usual spatiotemporal assignments that PL methods provide. We found that the approach is very versatile: TurboID can be used instead of APEX, and UV crosslinking can be used instead of FA crosslinking. The crosslinking step can even be skipped altogether and RBPs that have high RNA binding affinity will still be enriched. Moreover, APEX-PS may also be useful in other ways not explored here. The method should be straightforwardly extensible to other unpurifiable compartments, especially those already mapped by PL techniques including stress granules^60^, the ER membrane^45^, and lipid droplets^61^. In vivo organ-specific, cell-type specific, or organelle-specific RBP mapping should be possible, especially if APEX is replaced by TurboID; FA crosslinking of RNA to protein has been demonstrated in tissues^62^. In future work, it may be possible to develop variations of APEX-PS that provide RNA sequence information for the clients of APEX-biotinylated RBPs, or RNA binding site information for compartment-specific RBPs.

We used APEX-PS to investigate the biology of the outer mitochondrial membrane (OMM) and identified 13 RBPs at the OMM following PUR treatment. These proteins are candidates for recruiting and/or retaining the mitochondrial mRNAs that were observed at the OMM in our previous APEX-seq study ^24^. We examined one of our OMM RBP hits in-depth – SYNJ2BP – and discovered that it safeguards the OMM localization of its mRNA clients upon disassembly of polysomes and facilitates the translation of these mRNAs during mitochondrial recovery from stresses. As some OXPHOS-related mRNAs are clients of SYNJ2BP, we found that loss of SYNJ2BP impaired the activities of Complexes III and IV, as well as cell viability, during the stress recovery process. A working model is that SYNJ2BP binds to a subset of mitochondrial mRNAs and regulates their translation in response to inputs such as cell stress, ER status (due to SYNJ2BP’s role as a mitochondria-ER tether^45^), and transcriptional events.

## Figure legends

**Supplementary Figure 1.**
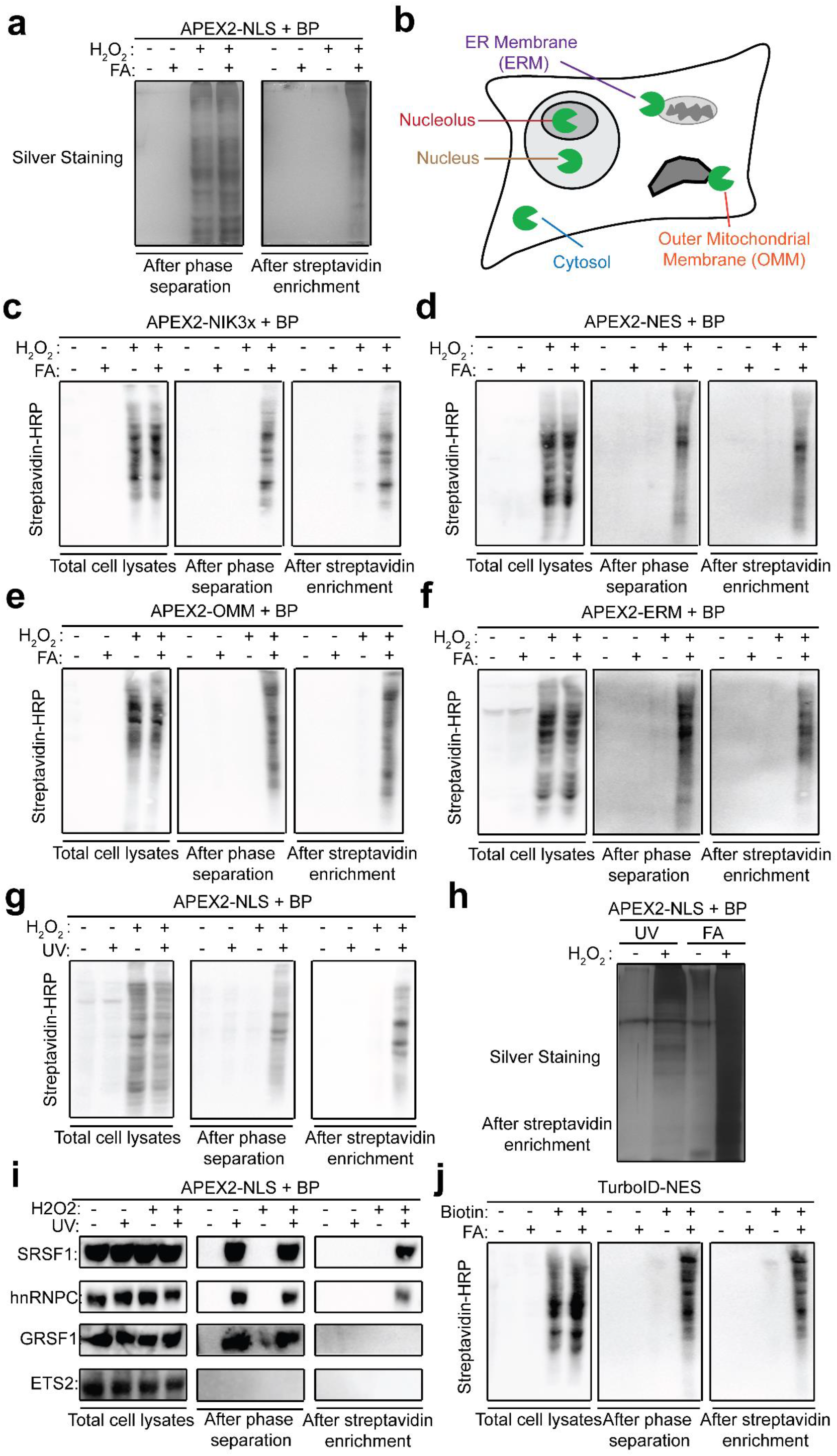
Evaluation of APEX-PS, related to Figure 1. **a**, Enrichment of subcellular RBPs. RBPs are enriched after phase separation. Biotinylated RBPs are enriched after the second enrichment using streptavidin beads. **b**, Schematic of five subcellular regions investigated in this study. **c-f**, Evaluation of APEX-PS in the nucleolus (**c**), cytosol (**d**), OMM (**e**) and ERM (**f**). **g**, UV crosslinking-based APEX-PS enables the enrichment of nuclear RBPs. After 1 minute of labeling with APEX-NLS, UV was applied to crosslink proteins to RNA at 254 nm with 400 mJ/cm^2^. Streptavidin blotting was performed after each step of the tandem enrichment. **h**, Comparison of FA- and UV-based APEX-PS. Silver staining of nuclear proteins was performed after streptavidin enrichment. **i**, After UV-based APEX-PS, we detected the four protein markers in Figure 1D using western blot. Samples blotted were whole cell lysate (left), material after phase separation (middle), and material after both phase separation and streptavidin bead enrichment (right). **j**, Capture of cytosolic RBPs by TurboID-PS. Cytosolic proteins were biotinylated by TurboID-NES and RNA-protein interactions were crosslinked by FA treatment. The samples were processed by phase separation and streptavidin enrichment, following by streptavidin blotting.

**Supplementary Figure 2.**
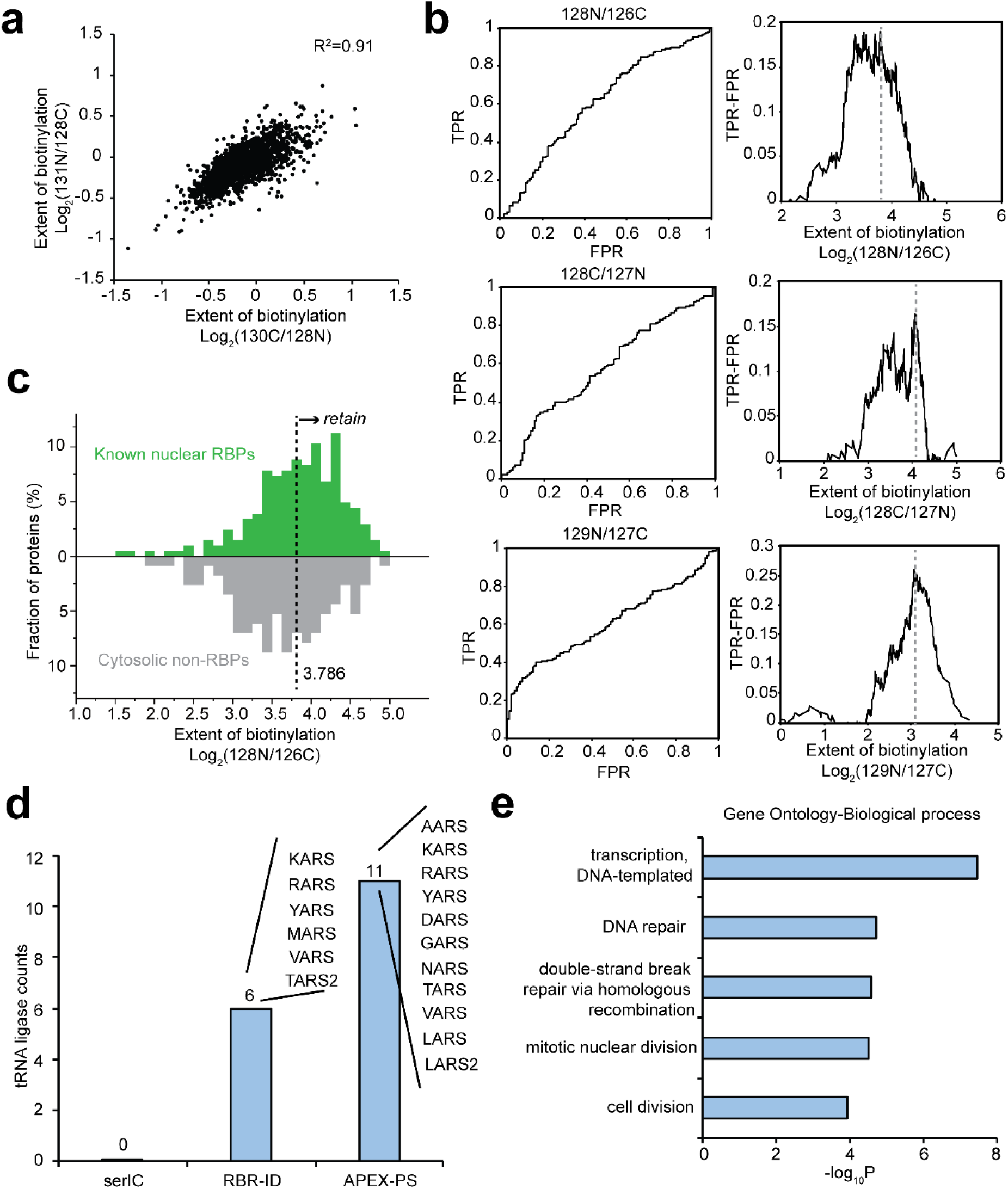
Additional analysis of nuclear APEX-PS dataset, related to Figure 2. **a**, Correlation of biological replicates. **b**, Receiver operating characteristic (ROC) curves of TMT ratios used for assignment of nuclear RBPs. Proteins were ranked in descending order based on the TMT ratio. True positive denotes known nuclear RBPs collected by overlapping known human RBPs and nuclear proteins annotated by GOCC (Table S1). False positives include cytosolic proteins that were not previously identified as RBPs. **c**, Sample histogram showing how the cutoff for 128N/126C ratio was applied. **d**, The number of tRNA ligases identified by APEX-PS and fractionation-based methods. The identified tRNA ligases were shown. (**E**) GO biological process analysis of nuclear RBP orphans identified by APEX-PS.

**Supplementary Figure 3.**
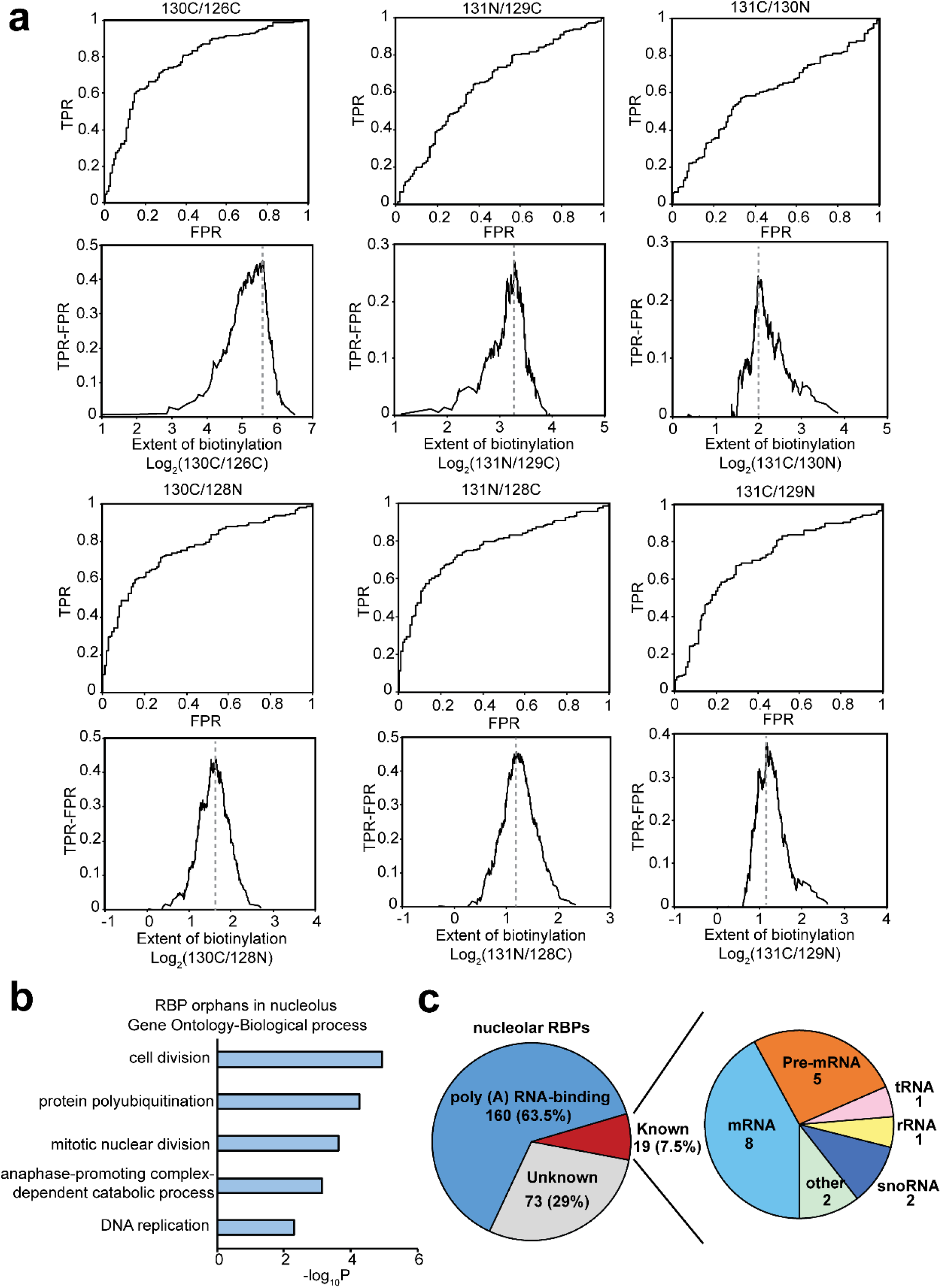
Additional analysis of nucleolar APEX-PS dataset, related to Figure 3. **a**, Receiver operating characteristic (ROC) curve of TMT ratios used for assignment of nucleolar RBPs. True positives are known nuclear RBPs collected by overlapping known human RBPs and nucleolar proteins annotated by GOCC (Table S1). False positives include cytosolic proteins that were not previously identified as RBPs. **b**, GO biological process analysis of nucleolar RBP orphans. **c**, Subclassification of RBPs in nucleolar APEX-PS dataset. Many RBPs we enriched bind to poly(A) ^6, 7, 12, 32, 34, 35^. Of the remainder, 19 have been experimentally shown to bind to the 6 RNA classes shown at right. The information of RNA types is shown in Supplementary Dataset 2.

**Supplementary Figure 4.**
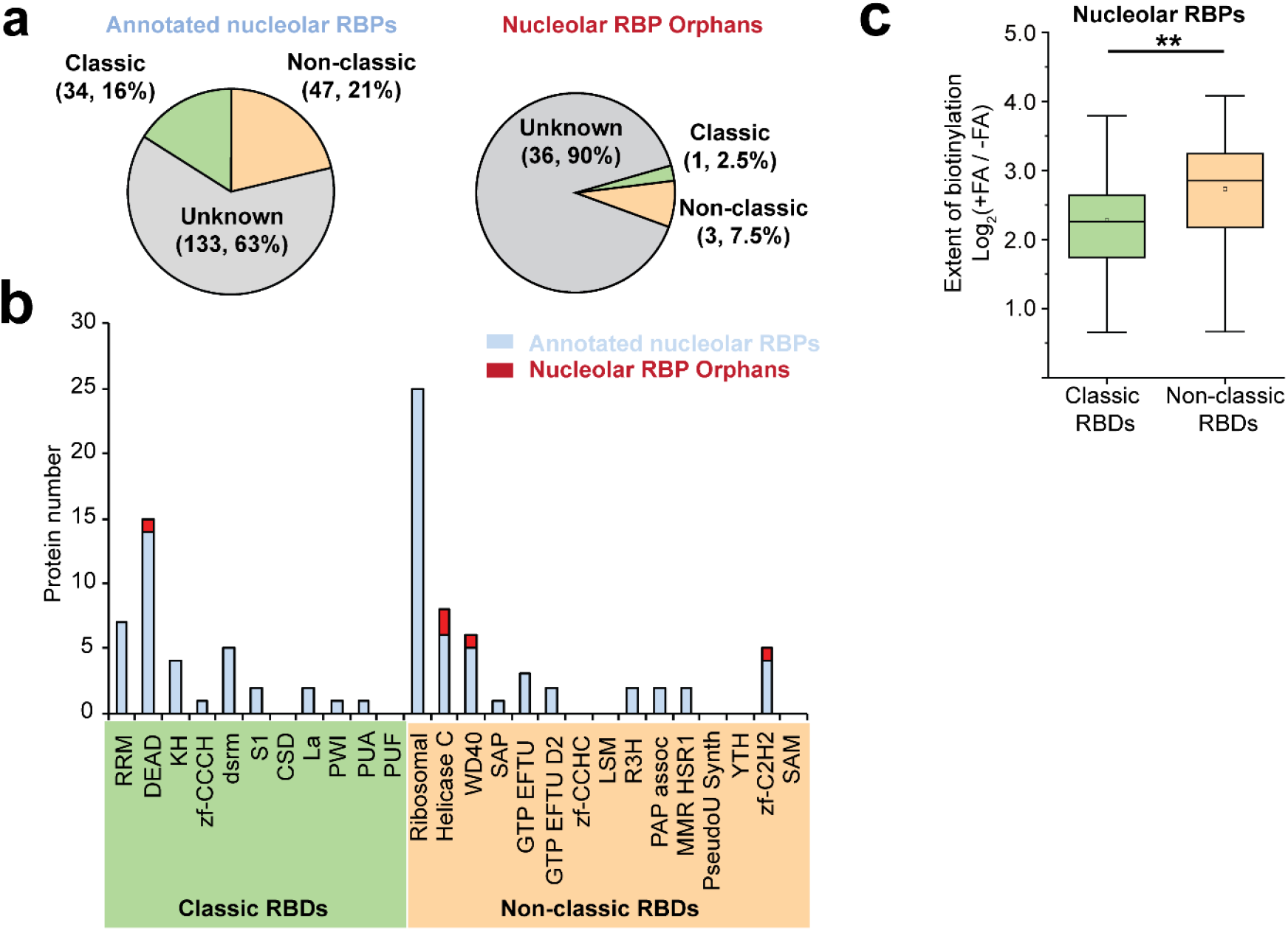
RNA binding domain (RBD) analysis for nucleolar APEX-PS dataset, related to Figure 4. **a**, Percentages of nucleolar RBPs with classical versus non-classical RNA binding domains (RBDs). **b**, The counts of RBPs with each type of classical and non-classical RNA binding domain. The RBDs of each RBP are listed in Table S2. **c**, Comparison of formaldehyde-dependent enrichment between classical (green) versus non-classical (yellow)-RBD-containing proteins in the nucleolar APEX-PS dataset. The relative biotinylation extent of +FA versus –FA is shown with the mean value of log_2_ (130C/130N), log_2_ (131N/130N) and log_2_ (131C/130N).

**Supplementary Figure 5.**
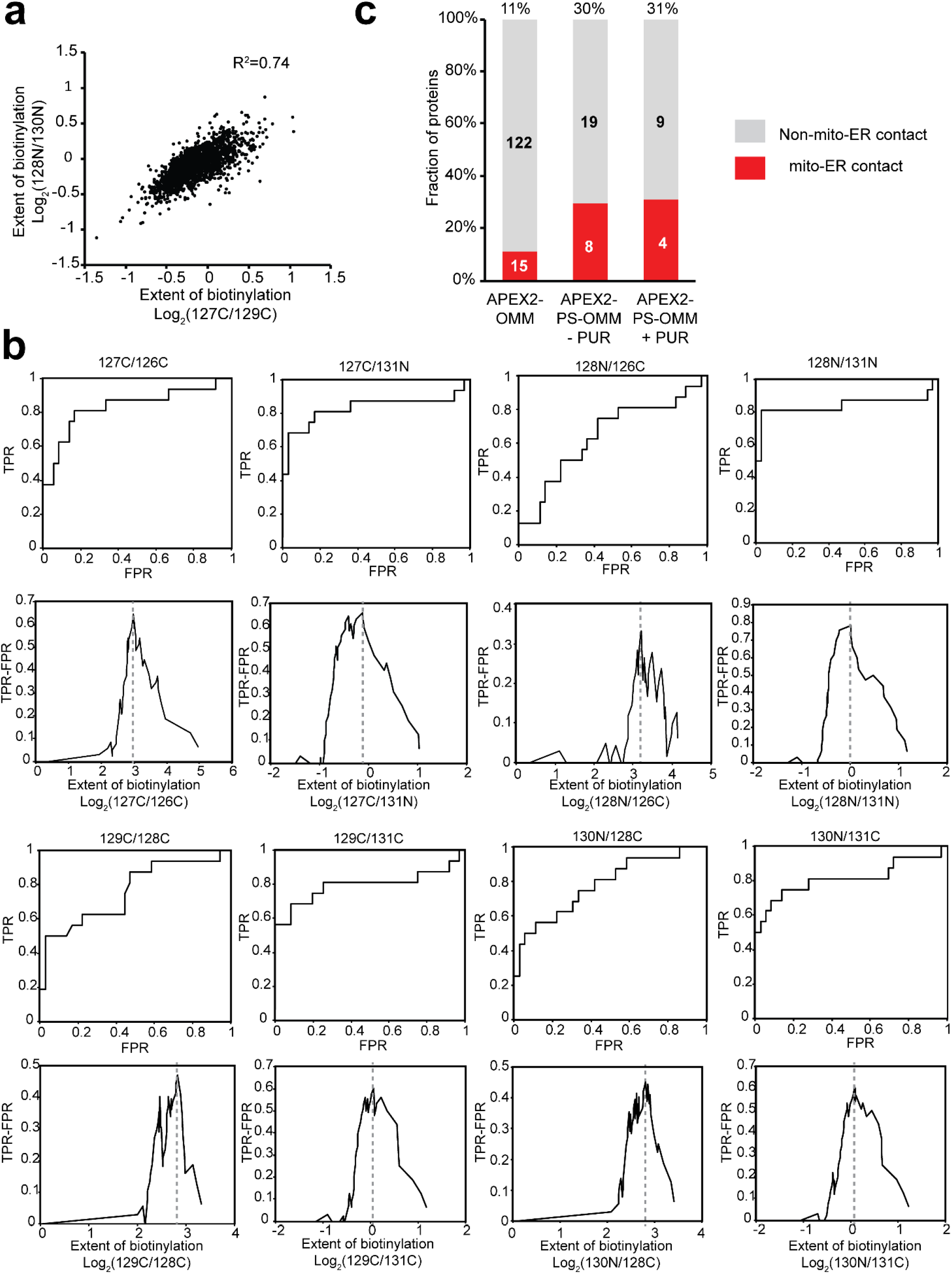
Additional analysis of mitochondrial APEX-PS dataset, related to Figure 5. **a**, Correlation between biological replicates for APEX2-PS-OMM profiling. **b**, Receiver operating characteristic (ROC) curve of TMT ratios used for assignment of OMM-localized RBPs. Proteins were ranked in descending order based on the TMT ratio. True positive denotes known OMM proteins annotated by GOCC. False positive includes cytosolic proteins that were not previously identified as RBPs. False positive proteins are mitochondrial matrix proteins identified by APEX profiling. **c**, Characterization of OMM-localized RBPs involved in mitochondrion-ER contact sites. The annotations of mitochondrial-ER contact are based on split-TurboID profiling^54^ and shown in Supplementary Table 4.

**Supplementary Figure 6.**
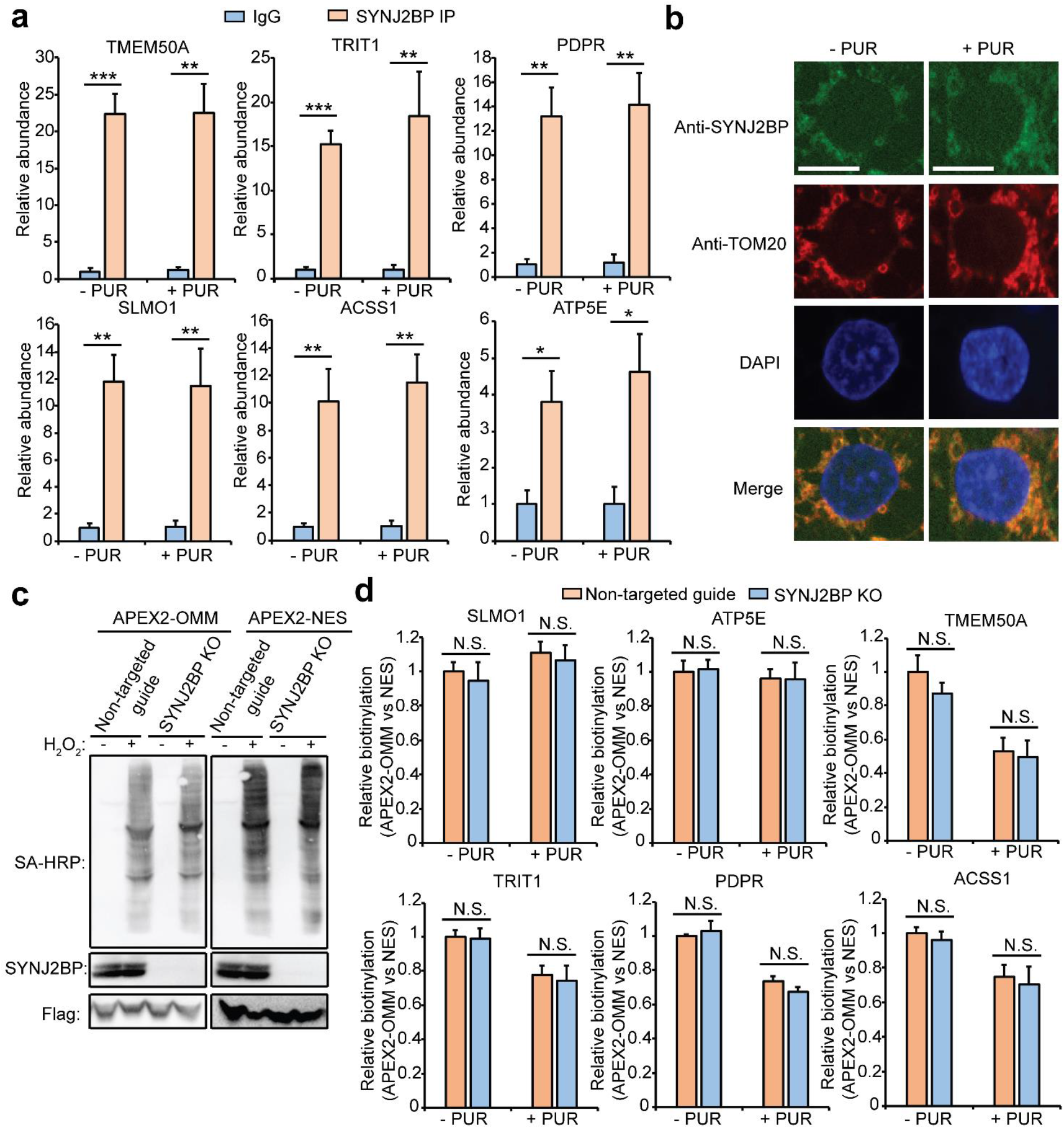
SYNJ2BP binds and promotes the OMM localization of its clients, related Figure 6. **a**, Validation of mitochondria-related SYNJ2BP clients in the absence or presence of protein translation inhibitor PUR by CLIP and qRT-PCR. **b**, Confocl imaging of SYNJ2BP localization at OMM under PUR condition. Anti-TOM20 stains OMM and DAPI stains nuclei. Scale bars, 10 μm. **c**, Validation of non-targeted guide control cells and SYNJ2BP knock-out cells stably expressing APEX2-OMM and APEX2-NES respectively. **d**, Evaluation of OMM localization of SYNJ2BP clients in SYNJ2BP knock-out cells by APEX-mediated proximity labeling of RNA.

**Supplementary Figure 7.**
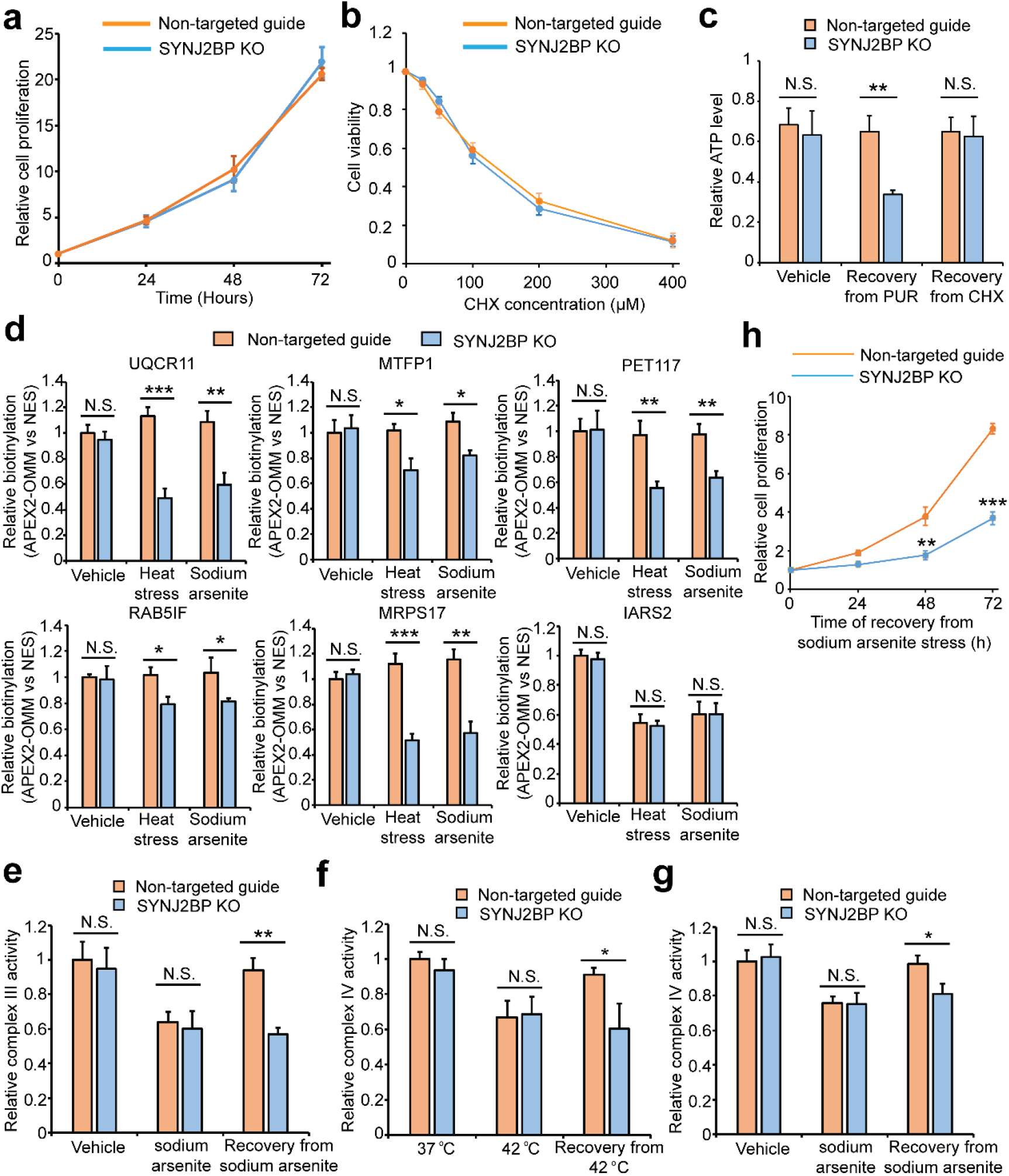
SYNJ2BP promotes the cellular recovery from stresses, related Figure 7. **a**, Evaluation of the cell viability of SYNJ2BP KO cells. **b**, Evaluation of cell viability of SYNJ2BP KO cells pre-treated with cycloheximide (CHX). **c**, Detection of ATP level in SYNJ2BP KO cells pre-treated with CHX and PUR. **d**, Evaluation of OMM localization of SYNJ2BP-regulated clients in SYNJ2BP knock-out cells by APEX-mediated proximity labeling of RNA. The OMM localization was determined by comparing APEX2-OMM with APEX2-NES labeling. IARS2 is an OMM-localized mRNA not identified to bind with SYNJ2BP. **e**, Evaluation of complex III activity in SYNJ2BP knock-out cells under sodium arsenite stress. **f-g**, Evaluation of complex IV activity in SYNJ2BP knock-out cells under heat (**f**) and sodium arsenite (**g**) stress. **h**, Evaluation of cell proliferation in SYNJ2BP knock-out cells during the recovery from sodium arsenite stress.

## Methods

### Cell Culture

HEK293T cells from the ATCC (passages <25) were cultured in a 1:1 DMEM/MEM mixture (Cellgro) supplemented with 10% fetal bovine serum, 100 units/mL penicillin, and 100 mg/mL streptomycin at 37°C under 5% CO_2_. For fluorescence microscopy imaging experiments, cells were grown on 7 × 7-mm glass coverslips in 48-well plates. For APEX-PS experiments, cells were grown on 15-cm glass-bottomed Petri dishes (Corning). To improve the adherence of HEK293T cells, glass slides and plates were pre-treated with 50 mg/mL fibronectin (Millipore) for 20 min at 37°C before cell plating and washing three times with Dulbecco’s PBS (DPBS) (pH 7.4). HEK293T cells stably expressing APEX2-NLS, APEX2-NIK3x, APEX2-OMM, APEX2-NES, ERM-APEX2 and TurboID-NES were generated in our previous studies ^17, 24^.

### APEX labeling and crosslinking in living cells

For both western blot analysis and proteomic analysis, HEK293T cells stably expressing the APEX2 fusion construct of interest were cultured in 15-cm dish for 18-24 h to about 90% confluency. APEX labeling was initiated by changing to fresh medium containing 500 μM biotin-phenol (Iris Biotech GMBH) and incubating at 37°C under 5% CO_2_ for 30 min. H_2_O_2_ (Sigma Aldrich) was then added to a final concentration of 1 mM and the plate was gently agitated for 1 min. For OMM-localized RBP profiling under PUR treatment, HEK293T cells stably expressing APEX2-OMM were treated with 200 μM puromycin and 500 μM biotin-phenol at 37°C under 5% CO_2_ for 30 min and then APEX labeling was initiated by H_2_O_2_ treatment for 1 minute.

For FA crosslinking, media was aspirated and the APEX labeling reaction was quenched by addition of 2 mL azide-free quenching solution (10 mM ascorbate and 5 mM Trolox in DPBS). Cells were incubated at room temperature for 1 min, then media was removed by aspiration and 10 mL of crosslink-quench solution (0.1% (v/v) formaldehyde, 10 mM sodium ascorbate and 5 mM Trolox in DPBS) was added. After 1 min, media were aspirated and cells were again incubated in 10 mL fresh crosslink-quench solution for 9 min at room temperature with gentle agitation. The crosslinking reaction was terminated in 125 mM of glycine for 5 min at room temperature. Cells were washed twice with 10 mL room-temperature DPBS, harvested by scraping, pelleted by centrifugation, and either processed immediately or flash frozen in liquid nitrogen and stored at -80°C for further analysis.

For UV crosslinking, the reaction was quenched by replacing the medium with an equal volume of quenching solution (10 mM ascorbate, 5 mM Trolox and 10mM sodium azide in DPBS). Cells were washed with quenching solution for three times and media were aspirated UV cross-linking was performed on PBS-washed cells by UV irradiation at 254 nm with 400 mJ/cm^2^ (CL-1000 Ultraviolet Crosslinker, UVP). Cells were washed twice with 10 mL ice-cold DPBS, harvested by scraping, pelleted by centrifugation, and either processed immediately or flash frozen in liquid nitrogen and stored at -80°C before further analysis.

For the validation of TurboID-PS shown in Figure S1J, HEK293T cells stably expressing TurboID-NES were cultured for 18-24 h. The biotin labeling was initiated by adding a final concentration of 50 μM biotin for 10 min. The labeling was stopped by transferring the cells to ice and washing five times with ice-cold DPBS. The cells were crosslinked by addition of 10 mL crosslinking solution (0.1% (v/v) formaldehyde in DPBS) for 10 min at room temperature with gentle agitation. The crosslinking reaction was terminated in 125 mM of glycine for 5 min at room temperature. Cells were washed and collected as described above.

### Phase separation

Cell pellets were resuspended in 1 mL Trizol and the homogenized lysate was transferred to a new tube. After incubating at room temperature for 5 min to dissociate, 200 μL of chloroform (Fisher Scientific) were added and vortexed, and the sample was centrifuged for 15 min at 12,000 g at 4 °C. The upper, aqueous phase (containing non-crosslinked RNAs) and the lower, organic phase (containing non-cross-linked proteins) was removed. Interface (containing the protein–RNA complexes) was resuspended in 1 mL Trizol and subjected to two more cycles of phase separation. Finally, the interface was precipitated by 1 mL methanol and pelleted by centrifugation at 14,000g, room temperature for 10 min. The precipitated proteins was resuspended in 100 μL of 100 mM triethylammonium bicarbonate (TEAB), 1 mM MgCl_2_, 1% SDS and incubated at 95°C for 20 min. The samples were cooled at room temperature for 5 min and digested with 2 μg RNase A, T1 mix (2 mg/mL of RNase A and 5,000 U/mL of RNase T1, Thermo Fisher Scientific) for 2 h at 37°C. Another 2 μg of RNase mix was added and incubated overnight at 37°C. The resulting solution was subjected to the final round of phase separation and the RBPs in organic phase was recovered by precipitation in 4.5 mL methanol with centrifugation at 14,000g, room temperature for 20 min. The protein pellets were resuspended in 1 mL methanol, transferred to a new 1.5 mL Eppendorf tube and pelleted by centrifugation at 14,000g, room temperature for 10 min. The protein pellets were resuspended in 1 mL RIPA buffer (50 mM Tris pH 8.0, 150 mM NaCl, 0.1% SDS, 0.5% sodium deoxycholate, 1% Triton X-100, 1× protease inhibitor cocktail (Sigma-Aldrich), and 1 mM PMSF) with sonication using a Misonix sonicator (0.5 s on, 0.5 s off, for a total of 10s on). The total RBP solution was then subjected for Western blotting or streptavidin enrichment.

### Streptavidin bead-based enrichment

To enrich biotinylated material from the total RBP solution, 300 μL streptavidin-coated magnetic beads (Pierce) were washed twice with RIPA buffer, then incubated with the 1 mL total RBP solution with rotation for 2 h at room temperature. The beads were subsequently washed twice with 1 mL of RIPA lysis buffer, once with 1 mL of 1 M KCl, once with 1 mL of 0.1 M Na_2_CO_3_, once with 1 mL of 2 M urea in 10 mM Tris-HCl (pH 8.0), and twice with 1 mL RIPA lysis buffer. For Western blotting analysis, the enriched proteins were eluted by boiling the beads in 75 μL of 3× protein loading buffer supplemented with 20 mM DTT and 2 mM biotin. For proteomic analysis, the beads were then resuspended in 1 mL fresh RIPA lysis buffer and transferred to a new Eppendorf tube. The beads were then washed with 1 mL wash buffer (75 mM NaCl in 50 mM Tris HCl pH 8.0) twice. The beads were resuspended in 50 μL of wash buffer and shipped to Steve Carr’s laboratory (Broad Institute) on dry ice for further processing and preparation for LC-MS/MS analysis.

### Gels and Western blots

For blotting shown in Figure 1C-D, Figure 6A and Figure S1, 5% of cells were subjected to total cell lysates analysis. the pellets were lysed by resuspending in 200 μL RIPA lysis buffer (50 mM Tris pH 8.0, 150 mM NaCl, 0.1% SDS, 0.5% sodium deoxycholate, 1% Triton X-100, 1× protease inhibitor cocktail (Sigma-Aldrich), and 1 mM PMSF) by gentle pipetting and incubating for 5 min at 4°C. Lysates were clarified by centrifugation at 10,000 r.p.m. for 10 min at 4°C. Protein concentration in clarified lysate was estimated with Pierce BCA Protein Assay Kit (ThermoFisher Scientific). 5% of total RBP solution was collected as the samples after phase separation. The total cell lysates, samples after phase separation and streptavidin enrichment were resolved on a 9% SDS-PAGE gel. The silver-stained gel shown in Figure S1A and H were generated using Pierce Silver Stain Kit (ThermoFisher Scientific).

For all Western blots, after SDS-PAGE, the gels were transferred to nitrocellulose membrane, and then stained by Ponceau S (5 min in 0.1% (w/v) Ponceau S in 5% acetic acid/water). The blots were then blocked in 5% (w/v) milk (LabScientific) in TBS-T (Tris-buffered saline, 0.1% Tween 20) for at least 30 min at room temperature. For streptavidin blotting, the blots were stained with 0.3 μg/mL streptavidin-HRP in 3% BSA (w/v) in TBS-T for 1 h at 4°C. The blots were washed three times with TBS-T for 5 min each time before to development. For validation of the specificity of APEX-PS in Figure 1D and S1I, the blots were stained with primary antibodies in 3% BSA (w/v) in TBS-T for 2 h in room temperature or overnight at 4°C. The primary antibodies include anti-SRSF1 (1:2500 dilution, ab129108, Abcam), anti-hnRNPC (1:2500 dilution, ab133607, abcam), anti-GRSF1 (1:2500 dilution, ab205531, Abcam) and anti-ETS-2 (1:1000 dilution, sc-365666, Santa Cruz Biotechnology). For validation of SYNJ2BP and EXD2 in Figure 6A and 6B, the blots were stained with anti-SYNJ2BP (1:2000 dilution, HPA000866-100UL, Sigma-Aldrich) and anti-EXD2 (1:2000 dilution, HPA005848-100UL, Sigma-Aldrich) in 3% BSA (w/v) in TBS-T for 2 h in room temperature or overnight at 4°C. For the evaluation of protein synthesis in Figure 6F, 6G, 7E and 7F, the blots were stained with anti-UQCR11 (1:1000 dilution, MBS715423, MyBioSource), anti-MTFP1 (1:2500 dilution, ab198217, Abcam), anti-PET117 (1:2500 dilution, PA5-61574, ThermoFisher Scientific), anti-RAB5IF (1:1000 dilution, PA543332, ThermoFisher Scientific), anti-MRPS17 (1:2500 dilution, ab175207, Abcam) and anti-HSP60 (1:2500 dilution, ab190828, Abcam) in 3% BSA (w/v) in TBS-T for 2 h in room temperature or overnight at 4°C. After washing three times with TBS-T for 5 min each, the blots were stained with secondary antibodies in 3% BSA (w/v) in TBS-T for 2 h in room temperature. The blots were washed three times with TBS-T for 5 min each time before to development with Clarity Western ECL Blotting Substrates (Bio-Rad) and imaging on the ChemiDoc XRS+ System (Bio-Rad).

### On-bead trypsin digestion of biotinylated proteins

To prepare samples for mass spectrometry analysis, proteins bound to streptavidin beads were washed twice with 200 μL of 50 mM Tris HCl buffer (pH 7.5) followed by two washes with 2 M urea/50 mM Tris (pH 7.5) buffer. The final volume of 2 M urea/50 mM Tris buffer (pH 7.5) was removed and beads were incubated with 50 μL of /50 mM Tris and 0.5 μg trypsin for 30 mins at 37°C with shaking. After 30 mins, the supernatant was removed and transferred to a fresh tube containing LysC digest. The streptavidin beads were washed once with 50 μL of 50 mM Tris buffer (pH 7.5) and the wash was combined with the on-bead digest supernatant and digested on shaker for at least 3 hours at 37 degrees. The eluate was reduced with 4 mM DTT for 30 min at 25°C with shaking. The samples were alkylated with 10 mM iodoacetamide for 45 min in the dark at 25°C with shaking. Then 0.5 μg of trypsin was added to the sample and the digestion continued overnight at 25°C with shaking. After digestion, samples were acidified (to pH < 3.0) by adding formic acid such that the sample contained ∼1% formic acid. Samples were desalted on C18 StageTips and evaporated to near dryness in a vacuum concentrator.

### TMT labeling and fractionation of peptides

Desalted peptides were labeled with TMT (11-plex) reagents ^63^. Peptides were reconstituted in 100 μL of 50 mM HEPES. Each 0.8 mg vial of TMT reagent was reconstituted in 41 μL of anhydrous acetonitrile and added to the corresponding peptide sample for 1 h at room temperature. Labeling of samples with TMT reagents was completed with the design shown in Figure 2A and Figure 5A. TMT labeling reactions were quenched with 8 μL of 5% hydroxylamine at room temperature for 15 min with shaking, evaporated to dryness in a vacuum concentrator, and desalted on C18 StageTips. For each TMT 11-plex cassette, 50% of the sample was fractionated by basic pH reversed phase using StageTips while the other 50% of each sample was reserved for LC-MS analysis by a single-shot, long gradient. One StageTip was prepared per sample using 2 plugs of Styrene Divinylbenzene (SDB) (3M) material. The StageTips were conditioned two times with 50 μL of 100% methanol, followed by 50 μL of 50% MeCN/0.1% formic acid, and two times with 75 μL of 0.1% formic acid. Sample was resuspended in 100 μL of 0.1% formic acid and loaded onto the stageTips and washed with 100 μL of 0.1% formic acid. Following this, sample was washed with 60 μL of 20 mM NH_4_HCO_3_/2% MeCN, this wash was saved and added to fraction 1. Next, sample was eluted from StageTip using the following concentrations of MeCN in 20 mM NH_4_HCO_3_: 10%, 15%, 20%, 25%, 30%, 40%, and 50%. For a total of 6 fractions, 10 and 40% (fractions 2 and 7) elutions were combined, as well as 15 and 50% elutions (fractions 3 and 8). The six fractions were dried by vacuum centrifugation.

#### Liquid chromatography and mass spectrometry

Fractionated peptides were resuspended in 8 μL of 0.1% formic acid and were analyzed by online nanoflow liquid chromatography tandem mass spectrometry (LC-MS/MS) using an Q-Exactive Plus Orbitrap MS (ThermoFisher Scientific) coupled online to an Easy-nLC 1200 (ThermoFisher Scientific). Four microliters of each sample was loaded onto a microcapillary column (360 μm outer diameter × 75 μm inner diameter) containing an integrated electrospray emitter tip (10 μm), packed to approximately 20 cm with ReproSil-Pur C18-AQ 1.9 μm beads (Dr. Maisch GmbH) and heated to 50 °C. The HPLC solvent A was 3% MeCN, 0.1% formic acid, and the solvent B was 90% MeCN, 0.1% formic acid. The SDB fractions were measured using a 110 min MS method, which used the following gradient profile at 200 nL/min: (min:%B) 0:2; 1:6; 85:30; 94:60; 95:90; 100:90; 101:50; 110:50 (the last two steps at 500 nL/min flow rate). The Q-Exactive Plus Orbitrap MS was operated in the data-dependent acquisition mode acquiring HCD MS/MS scans (resolution = 35,000, quadrupole isolation width of 0.7 Da) after each MS1 scan (resolution =70,000, 300-1800 m/z scan range) on the 12 most abundant ions using an MS1 target of 3×10^6^ and an MS2 target of 5×10^4^. The maximum ion time utilized for MS/MS scans was 120 ms and the HCD normalized collision energy was set to 30. The dynamic exclusion time was set to 20 s, and the peptide match and isotope exclusion functions were enabled. Charge exclusion was enabled for charge states that were unassigned, 1 and >6.

### Mass spectrometry data processing

Collected data were analyzed using Spectrum Mill software package v6.1pre-release (Agilent Technologies). Nearby MS scans with the similar precursor m/z were merged if they were within ± 60 s retention time and ±1.4 m/ z tolerance. MS/MS spectra were excluded from searching if they failed the quality filter by not having a precursor MH+ in the range of 750 - 4000. All extracted spectra were searched against a UniProt database (12/28/2017 containing human reference proteome sequences, common laboratory contaminants, and mycoplasma ribosomes. Search parameters included: parent and fragment mass tolerance of 20 p.p.m., 50% minimum matched peak intensity, and’calculate reversed database scores enabled. The digestion enzyme search parameter used was Trypsin Allow P, which allows K-P and R-P cleavages. The missed cleavage allowance was set to 3. TMT labeling was required at lysine, but peptide N termini were allowed to be either labeled or unlabeled. Allowed variable modifications were protein N-terminal acetylation, pyro-glutamic acid, deamidated N, and oxidized methionine. Individual spectra were automatically assigned a confidence score using the Spectrum Mill autovalidation module. Score at the peptide mode was based on target-decoy false discovery rate (FDR) of 1%. Protein polishing autovalidation was then applied using an auto thresholding strategy. Relative abundances of proteins were determined using TMT reporter ion intensity ratios from each MS/MS spectrum and the mean ratio is calculated from all MS/MS spectra contributing to a protein subgroup. Proteins identified by 2 or more distinct peptides and a protein score of at least 20 were considered for the dataset.

### Analysis of proteomic data for nucleus and nucleolus

To determine the cutoff in each biological replicate, we adopted the ratiometric analysis as previously described ^37^. The original identified proteins are shown in Table S1. For the assignment of nuclear RBPs, a list of known RBPs were firstly collected from RNA binding Gene Ontology Term (GO:0003723) and several previous datasets ^6, 7, 10, 12, 13, 32, 34, 35^. The true-positives (TPs) were the known RBPs with nuclear annotation in the following GO terms: GO:0016604, GO:0031965, GO:0016607, GO:0005730, GO:0001650, GO:0005654, GO:0005634. For the false-positives (FPs), a list of 6815 proteins with non-nuclear annotation in the following GO terms: GO:0015629, GO:0016235, GO:0030054, GO:0005813, GO:0045171, GO:0000932, GO:0005829, GO:0005783, GO:0005768, GO:0005929, GO:0005794, GO:0045111, GO:0005811, GO:0005764, GO:0005815, GO:0015630, GO:0030496, GO:0070938, GO:0005739, GO:0072686, GO:0005777, GO:0005886, GO:0043231; and are not annotated with the following GO terms: GO:0016604, GO:0031965, GO:0016607, GO:0005730, GO:0001650, GO:0005654, GO:0005634. The non-nuclear proteins that were not annotated as known RBP were assigned as FPs. For each replicate, the proteins were first ranked in a descending order according to the TMT ratio (128N/126C, 128C/127N, 129N/127N). For each protein on the ranked list, the accumulated true-positive count and false-positive count above its TMT ratio were calculated. A receiver operating characteristic (ROC) curve was plotted accordingly for each replicate (Figure S2B). The cutoff was set where true-positive rate - false-positive rate (TPR-FPR) maximized. Post-cutoff proteomic lists of the three biological replicates were intersected and proteins enriched in at least two biological replicates were collected. The potential glycosylated proteins were removed according to the annotation of glycoproteins or locations exclusively in the secretory pathway (e.g. ER/Golgi lumen, plasma membrane, extracellular regions) to obtain the nuclear RBPome list (Table S2).

For the assignment of nucleolar RBPs, TPs were known RBPs with nucleolar annotation (GO:0005730). The FPs were non-nuclear proteins without RBP annotation. For each replicate, the APEX-PS-NIK3x sample was not only compared to the negative controls (e.g. omitting H_2_O_2_ or enzyme), but also compared with the APEX-PS-NLS sample. The proteins were first ranked in a descending order according to the TMT ratio and cutoff was assigned by ROC analysis as described above (Figure S3A). The two types of comparison were intersected for each replicate (130C/126C and 130C/128N for replicate 1; 131N/129C and 131N/128C for replicate 2; 131C/129C and 131C/129N for replicate 3). The resulting lists of the three biological replicates were intersected and the potential glycosylated proteins were removed to obtain the final nucleolar RBP list (Table S3).

For the analysis of nuclear specificity of the nuclear and nucleolar RBPs (Figure 2H and 3F), we collected a list of 6889 human protein with nuclear annotations in the following GO terms: GO:0016604, GO:0031965, GO:0016607, GO:0005730, GO:0001650, GO:0005654, GO:0005634. The number of nuclear proteins presented in each dataset was determined. For the analysis of RNA binding specificity of nuclear RBPs (Figure 2G and 3G), the number of known RBPs presented in each dataset was determined. For the sensitivity analysis of nuclear RBPome (Figure 2I), a gold standard list of nuclear RBPs was manually curated (Table S2) according to previous literature^3^ and the coverage of APEX-PS, serIC and RBR-ID was determined. For comparing APEX-PS profiling with global phase separation profiling (Figure 2J and 3H), the protein abundance^36^ of overlapped RBPs and novel RBPs identified by APEX-PS was compared according to previous datasets. The analysis of the RNA types associated with nuclear and nucleolar RBPs (Figure 2F and S3C) was performed as previous studies^10^. Briefly, the RBPs identified by oligodT pulldown methods ^6, 7, 12, 32, 34, 35^ were assigned as poly(A) RNA binding proteins. The RNA binding types of the remaining RBPs were manually evaluated based on previous literature (Table S2 and S3). For the analysis of RBDs (Figure 4A, B and S4A, B), the domains of nuclear and nucleolar RBPs were obtained from Pfam (Table S2 and S3). The classification of classical and non-classical RBDs was based on previous studies ^7, 39^. The numbers of RBPs containing at least one classic RBD, only containing non-classical RBDs or without any RBDs were determined for both annotated RBPs and RBP orphans (Figure 4A and S4A). The number of nuclear and nucleolar RBPs containing each RBD was shown in Figure 4B and S4B, respectively. To identify RBPs with higher RNA binding activities, the nuclear RBPs were ranked in a descending order according to the +FA/-FA ratio (Mean value of 128N/127C, 128C/127C and 129N/127C for nuclear RBPs shown in Figure 4D). The ROC analysis was performed using RBPs with RRM as TPs and RBPs with non-classical RBDs as FPs. The nuclear RBPs with +FA/-FA ratio below the cutoff were assigned with high RNA binding affinity (Table S2).

### Analysis of proteomic data for OMM

The original identified proteins are shown in Table S4. To assign OMM RBPs under –PUR and +PUR conditions, a curated list of known OMM proteins^45^ was used as TPs and mitochondrial matrix proteins identified by APEX-mito profiling^16^ were assigned as FPs. For each replicate, the APEX-PS-OMM sample was not only compared to the negative control omitting H_2_O_2_, but also compared with the APEX-PS-NES sample. The proteins were first ranked in a descending order according to the TMT ratio and cutoff was assigned by ROC analysis as described above (Figure S5B). For assignment of OMM RBPs under basal condition, proteins above the cutoff of 127C/126C and 127C/131N were intersected for replicate 1 and proteins above the cutoff of 128N/126C and 128N/131N were intersected for replicate 2. For assignment of OMM RBPs under PUR treatment, proteins above the cutoff of 129C/128C and 129C/131C were intersected for replicate 1 and proteins above the cutoff of 130N/128C and 130N/131C were intersected for replicate 2. The resulting lists of the two biological replicates were intersected and the potential glycosylated proteins were removed to obtain the final OMM RBP list under basal and PUR condition, respectively (Table S5).

For the analysis of mitochondria specificity of the OMM RBPs (Figure 5E), a list of mitochondrial proteins were collected from MitoCarta database, GOCC terms containing mitochondrial annotations, mitochondrial matrix proteome and IMS proteome identified by APEX profiling. The number and percentage of mitochondrial proteins in human proteome, OMM proteins identified by APEX2-OMM profiling and the OMM RBPs under basal and PUR conditions was determined. For the analysis of OMM RBPs involved in mitochondrial-ER contact (Figure S5C), the number of OMM RBPs overlapped with proteins in mitochondrial-ER contact identified by split-TurboID^46^ was determined. For the analysis of RNA binding specificity of OMM RBPs (Figure 5F), the number of known RBPs described above in OMM RBPs was compared with that of OMM proteins identified by APEX profiling^45^.

### Immunofluorescence staining and fluorescence microscopy

For fluorescence imaging experiments in Figure 2B, HEK293T cells expressing APEX2-NLS and APEX2-NIK3x were plated and labeled as described above. Cells were fixed with 4% paraformaldehyde in PBS at room temperature for 15 min. Cells were then washed with PBS for three times and permeabilized with cold methanol at -20°C for 5-10 min. Cells were washed again three times with PBS and blocked for 1 h with 3% BSA in DPBS (‘‘blocking buffer’’) at room temperature. For APEX2-NLS imaging, cells were then incubated with anti-V5 antibody (1:1000 dilution, ThermoFisher Scientific) in blocking buffer for 1 h at room temperature. After washing three times with DPBS, cells were incubated with DAPI/secondary antibody (Alexa Fluor488), and neutravidin-Alexa Fluor647 in blocking buffer for 30 min. For APEX2-NIK3x imaging, cells were incubated with DAPI, and neutravidin-Alexa Fluor647 in blocking buffer for 30 min. Cells were then washed three times with DPBS and imaged. For fluorescence imaging in Figure S6B, HEK cells were treated with 200 μM puromycin for 30 min. Cells were fixed, washed and blocked as described above. Cells were incubated with anti-TOM20 (1:500 dilution, Santa Cruz Biotechnology) and anti-SYNJ2BP (1:500 dilution, Sigma-Aldrich) in blocking buffer for 1 h at room temperature.. After washing three times with DPBS, cells were incubated with DAPI/secondary antibody in blocking buffer for 30 min. Cells were then washed three times with DPBS and imaged. Fluorescence confocal microscopy was performed with a Zeiss AxioObserver microscope with 60× oil immersion objectives, outfitted with a Yokogawa spinning disk confocal head, Cascade II:512 camera, a Quad-band notch dichroic mirror (405/488/568/647), and 405 (diode), 491 (DPSS), 561 (DPSS) and 640 nm (diode) lasers (all 50 mW). DAPI (405 laser excitation, 445/40 emission), Alexa Fluor488 (491 laser excitation, 528/38 emission) and AlexaFluor647 (640 laser excitation, 700/75 emission) and differential interference contrast (DIC) images were acquired through a 60x oil-immersion lens. Acquisition times ranged from 100 to 2,000 ms. All images were collected and processed using SlideBook 6.0 software (Intelligent Imaging Innovations).

### Metabolic labeling of RNA-protein complexes

For the validation of SYNJ2BP and EXD2 as RBPs (Figure 6B), HEK293T cells were grown to ∼80% confluence in 15-cm dish and treated with 1 mM 5-EU for 16 h. The cells were washed with PBS for three times, followed by irradiation with 254-nm UV light at 150 mJ/cm^2^ (CL-1000 Ultraviolet Crosslinker, UVP). The cells were then lysed in 1 mL of 50 mM Tris-HCl (pH 7.5) buffer with sonication and subjected to centrifugation with 20000 *g* for 10 min to remove the debris. The lysates were reacted with 100 μM azide-PEG_3_-biotin (Click Chemistry Tools), 500 μM CuSO_4_, 2 mM THPTA (Sigma-Aldrich) and 5 mM sodium ascorbate (freshly prepared) for 2 h at r.t. with vortex, followed by adding 5 mM EDTA to stop the reaction. The lysates were precipitated with 8 vol of methanol at -80 °C for 1 h and washed twice with precooled methanol. The pellets were then resuspended in 1 mL RIPA lysis buffer with sonication and enriched by streptavidin beads overnight as we described above. After washing with RIPA buffer for three times, the beads were boiled in protein loading buffer with 2 mM biotin for 10 min. The samples were then analyzed by western blot with anti-SYNJ2BP and anti-EXD2 antibodies.

### RNA-immunoprecipitation sequencing (RIP-seq)

The RIP experiments were performed as described and high throughput sequencing services were provided by Cloud-Seq Biotech (Shanghai, China). The SYNJ2BP RIP was performed with SYNJ2BP antibody (Sigma Aldrich) and RNA-seq libraries were generated using the TruSeq Stranded Total RNA Library Prep Kit (Illumina) according to the manufacturer’s instructions and the library quality was evaluated with BioAnalyzer 2100 system (Agilent Technologies, Inc., USA). Library sequencing was performed on an illumina Hiseq instrument with 150 bp paired-end reads. The relative enrichment of each mRNA was obtained from the fold change of gene-level FPKM (fragments per kilobase of transcript per million mapped reads) values.

### Cross-linking immunoprecipitation (CLIP)

To validation of SYNJ2BP mRNA targets under – PUR and +PUR conditions, HEK293T cells were treated with 0 or 200 μM puromycin for 30 min. The cells were washed with 5 mL PBS for three times, crosslinked by 254-nm UV light at 150 mJ/cm^2^. Then the cells were lysed in 500 μL CLIP lysis buffer (50 mM Tris·HCl, pH 7.5, 150 mM NaCl, 1% Nonidet P-40, 0.1% SDS, and EDTA-free protease inhibitor mixture). The cell lysates were incubated on ice for 10 min and cleared by centrifugation at 13000 *g* for 15 min at 4 °C. 125 μL Dynabeads protein G (ThermoFisher Scientific) were washed with 500 μL lysis buffer for twice and incubated with 10 μg anti-SYNJ2BP antibody at r.t. for 45 min. The lysates were then incubated with the antibody-conjugated beads overnight at 4 °C. The beads were washed twice with 900 μL high salt wash buffer (50 mM Tris-HCl pH 7.4, 1 M NaCl, 1 mM EDTA, 1% NP-40, 0.1% SDS and 0.5% sodium deoxycholate) and twice with 500 μL wash buffer (20 mM Tris-HCl pH 7.4, 10 mM MgCl_2_, 0.2% Tween-20). The beads were resuspended in 54 μL water, and then mixed with 33 μL 3X proteinase digestion buffer, 10 μL proteinase K (20 mg/mL, ThermoFisher Scientifi) and 3 μL Ribolock RNase inhibitor. The 3X proteinase digestion buffer were freshly prepared as follow: 330 μL 10X PBS, pH = 7.4 (Ambion); 330 μL 20% N-laurylsarcosine sodium solution (Sigma-Aldrich); 66 μL of 0.5 M ETDA; 16.5 μL of 1 M DTT; 357.5 μL water. Proteinase digestion was performed at 42 °C for 1 h and 55 °C for 1 h with vigorous mixing and the supernatant was collected. The recovered RNAs were purified using RNA clean and concentrator -5 kit (Zymo Research) and subjected to further analysis.

### Generation of SYNJ2BP KO cells stably expressing APEX2 constructs

The non-targeted guide and SYNJ2BP KO HEK293T cells were generated previously ^45^. For preparation of lentiviruses, HEK293T cells in 6-well plates were transfected at 60%–70% confluency with the lentiviral vector pLX304 containing APEX2-OMM or APEX-NES (1,000 ng), the lentiviral packaging plasmids dR8.91 (900 ng) and pVSV-G (100 ng), and 8 mL of Lipofectamine 2000 for 4 h. About 48 h after transfection the cell medium containing lentivirus was harvested and filtered through a 0.45-mm filter. The non-targeted guide and SYNJ2BP KO HEK293T cells were then infected at ∼50% confluency, followed by selection with 8 mg/mL blasticidin in growth medium for 7 days before further analysis.

### APEX RNA labeling at OMM

APEX labeling was performed as described above in non-targeted guide and SYNJ2BP KO HEK293T cells stably expressing APEX2-OMM or APEX2-NES. The RNA was extracted from cells using the RNeasy plus mini kit (QIAGEN) following the manufacture protocol, including adding β-mercaptoethanol to the lysis buffer. The cells were sent through the genomic DNA (gDNA) eliminator column supplied with the kit. A modification to the protocol was replacing the RW1 buffer with RWT buffer (QIAGEN) for washing. The extracted RNA was eluted into RNase-free water and RNA concentrations were determined using the Nanodrop (ThermoFischer Scientific).

To enrich biotinylated RNAs, we used 10 μL Pierce streptavidin magnetic beads (ThermoFischer Scientific) per 25 mg of RNA. The beads were washed 3 times in B&W buffer (5 mM Tris-HCl, pH 7.5, 0.5 mM EDTA, 1 M NaCl, 0.1% TWEEN 20), followed by 2 times in Solution A (0.1 M NaOH and 0.05 M NaCl), and 1 time in Solution B (0.1 M NaCl). The beads were then suspended in 100-150 mL 0.1 M NaCl and incubated with 100-125 mL RNA on a rotator for 2 h at 4°C. The beads were then placed on a magnet and the supernatant discarded. Beads were washed 3 times in B&W buffer and resuspended in 54 mL water. A 3X proteinase digestion buffer was made (1.1 mL buffer contained 330 mL 10X PBS pH = 7.4 (Ambion), 330 μL 20% N-Lauryl sarcosine sodium solution (Sigma Aldrich), 66 mL 0.5M EDTA, 16.5 mL 1Mdithiothreitol (DTT, ThermoFischer Scientific) and 357.5 mL water). 33 μL of this 3X proteinase buffer was added to the beads along with 10 mL Proteinase K (20 mg/mL, Ambion) and 3 mL Ribolock RNase inhibitor. The beads were then incubated at 42°C for 1 h, followed by 55°C for 1 h on a shaker. The RNA was then purified using the RNA clean and concentrator -5 kit (Zymo Research) and subjected to further analysis.

### RT-Qpcr

For the RT-qPCR analysis of CLIP and APEX RNA labeling experiments, the enriched RNA was first reverse transcribed following the Superscript III reverse transcriptase (ThermoFischer Scientific) protocol using random hexamers as primers. The resulting cDNA was then tested using qPCR using the primers above in 2X SYBR Green PCR Master Mix (ThermoFischer Scientific), with data generated on Lightcycler 480 (Roche).

### Azidohomoalanine labeling

To evaluate the impact of SYNJ2BP on protein synthesis of its clients (Figure 6G, 7E and 7F), cells were cultured in methionine-free medium supplemented with 1 mM azidohomoalanine (AHA). Cells were lysed in RIPA buffer and protein concentration was normalized to 2 mg/mL. 1 mL lysates were reacted with 100 μM biotin-PEG4-alkyne, premixed 2-(4-((bis((1-tertbutyl-1H-1,2,3-triazol-4-yl)methyl)amino)methyl)-1H-1,2,3-triazol1-yl)-acetic acid (BTTAA)-CuSO4 complex (500 μM CuSO4, BTTAA:CuSO4 with a 2:1 molar ratio) and 2.5 mM freshly prepared sodium ascorbate for 2 h at room temperature. The resulting lysates were precipitated by 8 mL methanol at -80°C overnight and the precipitated proteins were centrifuged at 8000 *g* for 5 min at 4 °C. The proteins were washed twice with 1 mL cold methanol and resuspended in 1 mL RIPA buffer with sonication. The biotinylated proteins were further captured by 200 μL streptavidin magnetic beads for 2 h. The beads were washed as described above and proteins were eluted by boiling the beads in 75 μL of 3× protein loading buffer supplemented with 20 mM DTT and 2 mM biotin. The resulting samples were analyzed by western bloting with antibodies indicated.

### Complex III and IV activity assay

Complex III activity was assayed using a mitochondrial complex III activity assay kit (Sigma Aldrich) and complex IV activity was determined using a complex IV human enzyme activity microplate assay kit (Abcam). HEK293T expressing non-targeted Cas9 and SYNJ2BP knockout cells were obtained from our previous study ^45^ were plated in 15 cm dish. Cell pellets were lysed and mitochondrion was purified according to the manufacturer’s protocol in a mitochondrial isolation kit for cultured cells (Abcam). The activity was determined by following the manufacturer’s protocol with a standard curve.

### Cell proliferation assays

In order to determine the effect of SYNJ2BP on cell proliferation (Figure S7A), HEK293T expressing non-targeted Cas9 and SYNJ2BP knockout cells were obtained from our previous study ^45^. The MTS assay was performed using the CellTiter 96 AQueous One Solution Cell Proliferation Assay kit (Promega), following the manufacturer’s instructions. 1 × 10^4^ cells per well were plated in 96-well plates with 100 μL fresh medium per well. The cells were cultured for 1-3 days, and the medium was freshly changed every 24 h. 20 μL of CellTiter 96 AQueous One Solution Reagent was added into each well and incubated for 4 h.

To evaluate the impact of SYNJ2BP knockout on cell viability under CHX and PUR treatment (Figure 7A and S7B), 1 × 10^4^ cells per well were plated in 96-well plates with 100 μL fresh medium per well. After 24 h, the cells were treated with desired concentration (0, 25, 50, 100, 200 and 400 μM) of drugs for 12 h and then changed into normal medium for another 12 h. For glucose/galactose cell viability assay (Figure 7B), 2000 cells per well were plated in 96-well plates with 100 μL fresh medium per well. After 24 h, cells were washed with DPBS and the growth medium was replaced with medium containing 10% FBS, 100 units/mL penicillin, 100 mg/mL streptomycin and DMEM without glucose supplemented with 10 mM galactose or 10 mM glucose, as well as 200 μM drugs. After 24 h, the cells were changed into the mediums without drugs for 48 h and 20 μL of CellTiter 96 AQueous One Solution Reagent was added into each well and incubated for 4 h.

To evaluate the cellular recovery from heat stress (Figure 7H), 1 × 10^4^ cells per well were plated in 96-well plates with 100 μL fresh medium per well. After 24 h, the cells were incubated at 42°C for 1 h and then incubated at 37°C for 1-3 days. For the sodium arsenite stress, cells were treated with 400 μM sodium arsenite for 1 h and then cultured in the normal medium for 1-3 days. 20 μL of CellTiter 96 AQueous One Solution Reagent was added into each well and incubated for 4 h. The absorbance at 490 nm was recorded using a 96-well plate reader. Each biological experiment has five technical replicates and three biological replicates were performed.

### Statistical analysis

For comparison between two groups, P values were determined using two-tailed Student’s t tests, *P < 0.05; **P < 0.01; ***P < 0.001; N.S. not significant. For all box plots (Figure 3c, Supplementary Figure 4c, 5c, 5d and 6d), P values were calculated with Wilcoxon rank sum by R (*P < 0.05; **P < 0.01; ***P < 0.001). Error bars represent means ± SD.

### Reporting Summary

Further information on research design is available in the Nature Research Reporting Summary linked to this article.

### Data availability

Additional data beyond that provided in the Figures and Supplementary Information are available from the corresponding author upon request.

## Acknowledgements

We are grateful to the NIH (U24-CA210986 to S.A.C., U01-CA214125 to S.A.C., R01-DK121409 to S.A.C. and A.Y.T.), Chan Zuckerberg Biohub, and Beckman Technology Development Seed Grant for support of this work. S. Han (Stanford University) provided images for APEX2-NLS and APEX2-NIK3x labeling.

## Author contributions

W.Q. and A.Y.T. designed the research and analyzed all the data except where noted. W.Q. performed all experiments except where noted. W.Q., A.Y.T., S.A.M., and S.A.C. designed the proteomics experiments. W.Q. prepared the proteomic samples. S.A.M. and D.K.C. processed the proteomic samples and analyzed the MS data. W.Q. and A.Y.T. wrote the paper with input from all authors.

